# *SORBS2* is a genetic factor contributing to cardiac malformation of 4q deletion syndrome

**DOI:** 10.1101/2020.05.12.087452

**Authors:** Fei Liang, Xiaoqing Zhang, Bo Wang, Juan Geng, Guoling You, Jingjing Fa, Huiying Sun, Huiwen Chen, Qihua Fu, Zhen Zhang

## Abstract

Chromosome 4q deletion is one of the most frequently detected genomic imbalance events in congenital heart disease (CHD) patients. However, a portion of CHD-associated 4q deletions do not include known CHD genes. Alignment of those 4q deletions defined a minimal overlapping region including only one gene-*SORBS2*. Histological analysis of *Sorbs2*^*-/-*^ heart revealed atrial septal hypoplasia/aplasia or double atrial septum. Mechanistically, *SORBS2* had a dual role in maintaining sarcomeric integrity of cardiomyocytes and specifying the fate of second heart field (SHF) progenitors through c-ABL/NOTCH/SHH axis. In a targeted sequencing of a panel of known and candidate CHD genes on 300 CHD cases, we found that rare *SORBS2* variants were significantly enriched in CHD patients. Our findings indicate that *SORBS2* is a regulator of cardiac development and its haploinsufficiency may contribute to cardiac phenotype of 4q deletion syndrome. The presence of double atrial septum in *Sorbs2*^*-/-*^ hearts reveals the first molecular etiology of this rare anomaly linked to paradoxical thromboembolism.

## Introduction

Congenital heart disease (CHD) is present in ∼1% newborns (1). Although CHD is the most common birth defect, the underlying causes of most CHDs remain elusive. Genetic heterogeneity and gene-environment interaction limit the genotype-phenotype correlation and complicate interpretation of CHD genetics. Anatomic rectification through interventional and surgical treatments have greatly improved survival rate of CHD patients (2). However, the same medical treatment often yields different outcomes in different CHD patients, suggesting the underlying genetic lesion impacts CHD prognosis. Therefore, elucidation of CHD genetic basis is becoming ever more important in both reducing short-term therapeutic complications and improving long-term care of CHD patients. With the advances in genetic diagnosis techniques, the discovery of genetic variants is no longer a limit (3). The interpretation of the association of identified genetic variants with phenotype becomes the imperative need to address the genetic basis of CHD.

CNV is a chromosomal structural aberration comprising of deletion or duplication with a size larger than 1kb. Due to the abundance of low-copy repeats and retrotransposons, CNVs are quite common genetic variants in human genomes. Previous studies have estimated that CNVs contribute to 10-15% CHD (4, 5). The most common chromosomal deletion is Del22q11.2, which is the shared genetic cause of 3 clinical syndromes—DiGeorge syndrome, velocardiofacial syndrome and conotruncal anomaly face syndrome (6). The common forms of Del22q11.2 are ∼3 or 1.5 Mbs deletion encompassing dozens of genes (6). The successful identification of the major CHD gene *TBX1* has greatly advanced our understanding of the pathogenesis of 22q11.2 deletion syndrome (22q11.2DS) (7-10). The gained knowledge also leads to the discovery of potential therapeutic measures to rescue cardiovascular defect through pharmaceutical manipulation of *TBX1*-involved biological pathway (11-13). Interestingly, untypical 22q11.2 deletions outside of the 1.5 Mbs critical region are also associated with CHD(14-16). Studies have shown that *CRKL* and *LZTR1* within these untypical deletions regulate cardiac development and may contribute to heart malformations of patients with typical 3Mbs deletion(15, 17, 18). Therefore, the identification of all the disease genes within CNV intervals is crucial for having a complete view of the pathogenesis of related phenotypes.

We previously profiled CNVs distribution in 514 CHD cases and have found that chromosome 4q deletion is the most frequent CHD-related CNV in our cohort(19). The majority of terminal 4q deletion we identified does not include *HAND2* that are considered as the culprit gene responsible for CHD in 4q deletion syndrome. Through comparing our findings with reported 4q deletions, we hypothesized that *SORBS2* within the minimal overlapping region may be a CHD gene. To this end, we deleted *Sorbs2* in mice. Histological analyses of *Sorbs2*^*-/-*^ mouse hearts revealed atrial septal anomalies in 40% (12/30) mice. *SORBS2* knockdown in *in vitro* human cardiomyocyte differentiation model indicated a dual role of *SORBS2* in cardiogenesis. The encoded protein not only maintains the sarcomeric integrity as a structural component, but also promotes the differentiation of the second heart field (SHF) progenitors through C-abl/Notch1/Shh axis. To further validate its pathogenicity in CHD, we performed targeted sequencing of 104 candidate and known CHD genes on 300 complex CHD cases. After comparing with 220 Han Chinese subjects from 1000 genomes, we found that *SORBS2* and known CHD gene *KMT2D* had significantly higher mutational burden in the CHD group. Taken together, our data indicate that *SORBS2* is a novel CHD gene contributing to the cardiac phenotype of 4q deletion syndrome.

## Results

### *SORBS2* is a potential CHD disease gene

We aligned CHD-associated terminal 4q deletions identified in our previous study with reported 4q deletions. We noted that more than half of them do not include the known CHD gene *HAND2* (Figure 1A). The minimal overlapping region among those deletions includes only one gene-*SORBS2*. It was proposed as a candidate CHD gene in the study identifying this unusual small deletion(20). However, the pathogenicity of *SORBS2* deficiency on CHD is still unclear.

**Figure 1.**
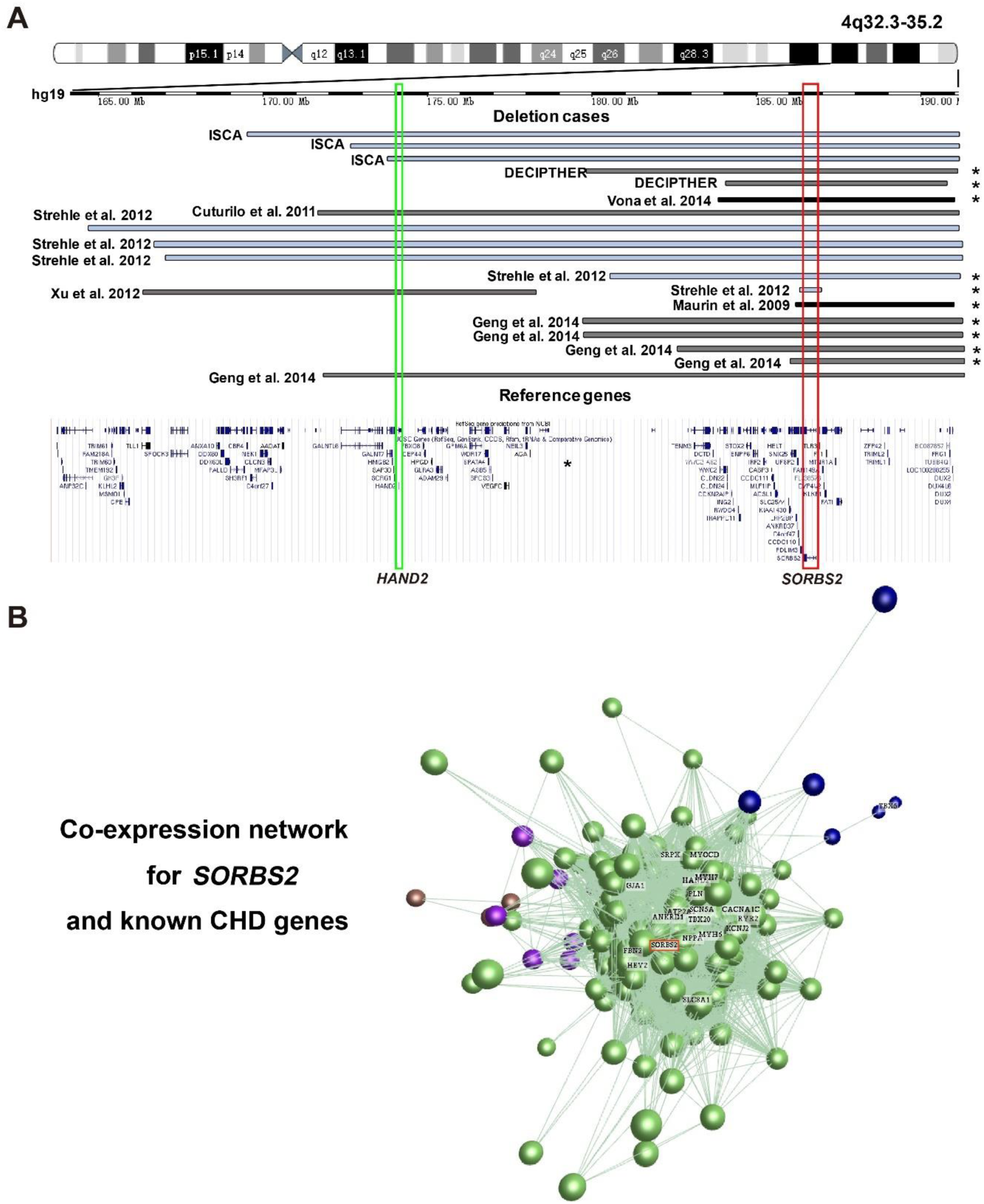
*SORBS2* is a candidate CHD gene. A. Alignment of 4q deletions identified by us and others(19, 37, 38, 40, 63, 64). Asterisk, 4q deletions not including *HAND2*, the known cardiac development regulator responsible for CHD of 4q deletion syndrome. Green box, the *HAND2* gene. Red box, the *SORBS2* gene within the minimal overlapping region among 4q deletions indicated by asterisks. B. Co-regulation network of *SORBS2* with multiple known CHD-related genes. *SORBS2* is highlighted in red box. Color-coded dots represent different clusters.

*SORBS2* encodes a cytoskeleton associated adaptor protein that regulates cytoskeletal organization and intracellular signaling (21). It is highly expressed in the heart and participates in structural organization of cardiomyocyte Z-discs (22). In analyzing RNA-seq data of human fetal heart tissues (96 days -147 days) from the ENCODE database (23), we found that *SORBS2* expression was highly correlated with those of multiple known CHD genes (Figure 1B). Taken together, we thought that *SORBS2* might be required for cardiac development.

### *Sorbs2*^*-/-*^ mice have atrial septal defect

The entire *SORBS2* gene is absent in terminal 4q deletion, hence we used *Sorbs2* knockout mice to study its role in cardiac development. In previous report, about 40-60% *Sorbs2*^*-/-*^ mice die within 1 week after birth (24). We wonder whether structural heart defect(s) contributes to the early lethality. To this end, we intercrossed *Sorbs2*^*+/*-^ mice and collected 137 embryos at E18.5. The ratio of genotype distribution among embryos is consistent with Mendel’s law (Table S1), suggesting no embryo loss in early development stage. We dissected 30 *Sorbs2*^*-/-*^ embryos and none of them showed conotruncal defect from gross examination (Figure 2A). Next, we performed paraffin sectioning for intracardiac morphological analysis. No ventricular septal defect (VSD) was noted in all the *Sorbs2*^*-/-*^ hearts (Figure 2B). However, we found that about 40% (12/30) *Sorbs2*^*-/-*^ hearts had atrial septal defect (ASD) with 10 being the absence of primary septum and 2 being DAS (Figure 2C). The presence of ASD in *Sorbs2*^*-/-*^ mice indicates that *SORBS2* deficiency contributes to CHD pathogenesis.

**Figure 2.**
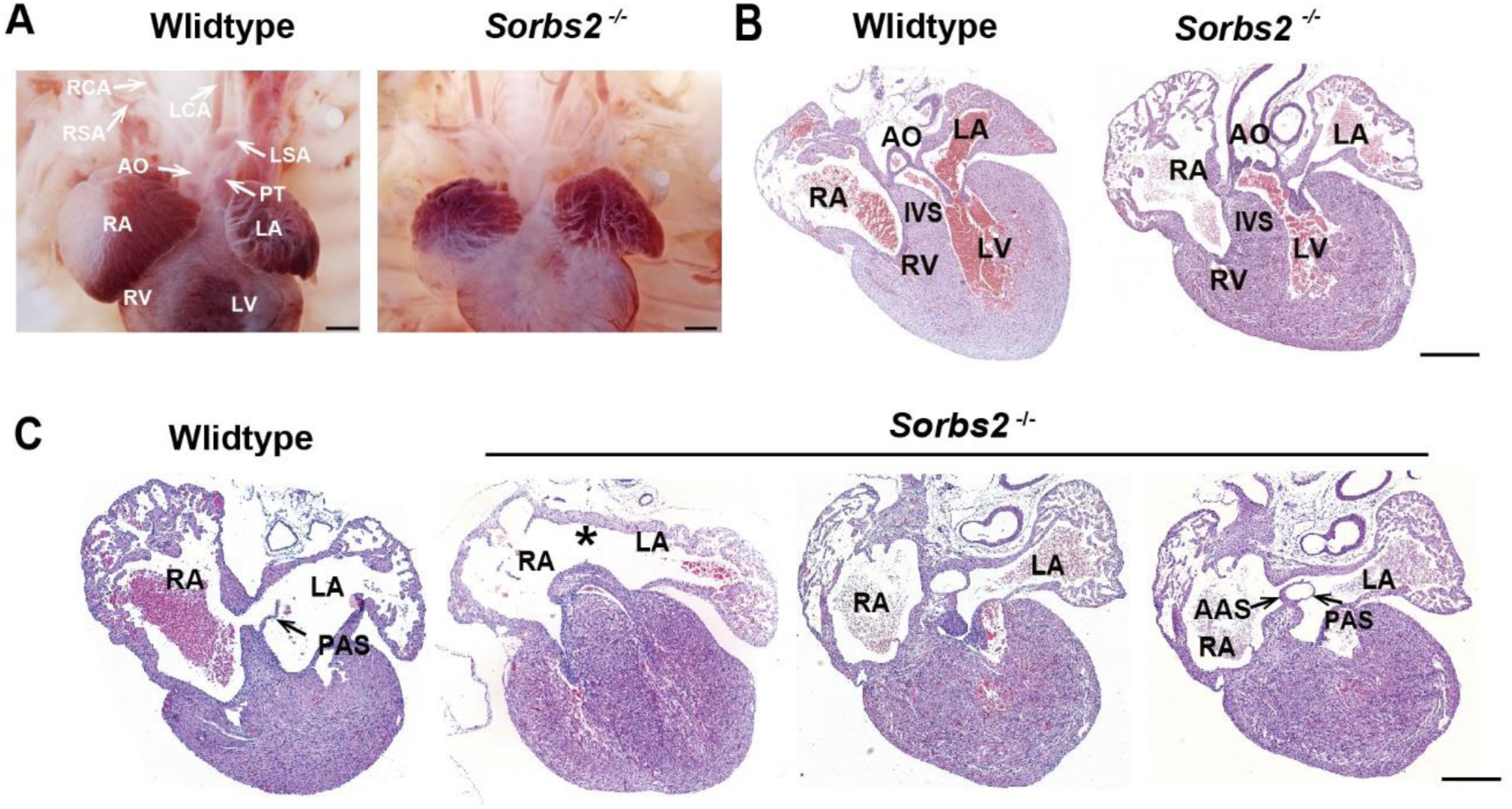
Cardiac phenotype of *Sorbs2*^*-/-*^ mice. A. Gross view of embryos at E18.5. B. HE-stained paraffin sections of E18.5 hearts in conotruncal area. C. HE-stained paraffin sections of E18.5 heart in atrial septum area. Asterisk indicates the absence of PAS. Two sections in the right are from the same heart with DAS. The rightmost section is dorsal to the other. AO, aorta. PT, pulmonary trunk. LSA, left subclavian artery. RSA, right subclavian artery. LCA, left common carotid artery. RCA, right common carotid artery. LA, left atrium. RA, right atrium. LV, left ventricle. RV, right ventricle. PAS, primary atrial septum. AAS, accessory atrial septum. IVS, interventricular septum. Scale bar: 1mm in A, 200μm in B and C.

### *SORBS2*-knockdown cardiomyocytes have disrupted sarcomeric structure but no obvious electrophysiological abnormality

Having known that *Sorbs2* knockout caused ASD in mice, we used in vitro stem cell-based cardiogenesis to study whether *SORBS2* has a conserved role in human. Since 4q deletion causes *SORBS2* haploinsufficiency, we approached the question by knocking down *SORBS2* in H1 human ES cell lines (H1-hESC). We used two different shRNAs to knock down *SORBS2* and similar knockdown efficiencies (∼40% of wild type expression level) were achieved (Figure S1A). Compared to scramble-shRNA controls, there was no significant difference in the growth rate and clone morphology of *SORBS2*-knockdown hESCs. To verify this observation, we examined the expression of pluripotency markers Oct4, Sox2, Nanog, TRA-1-60, and no significant difference was detected on both RNA and protein levels between control and *SORBS2*-knockdown groups (Figure S1B-S1E). We used a well-established 2-dimensional monolayer differentiation protocol to perform cardiomyocyte differentiation (25) (Figure S2A). Since *SORBS2*-knockdown ES cell lines from different shRNAs had similar phenotypes (Figure S2B, Video S1-S3), we only used *SORBS2-shRNA1* for further analyses. Spontaneous beating started to appear at differentiation D8 and robust contraction occurred at D10 in control hESCs. Although spontaneous beating also appeared at D8 in *SORBS2*-knockdown group, differentiated cardiomyocytes contracted much weaker (Video EV1-EV3). Since SORBS2 is a structural component of sarcomeric Z-line (26), we examined the myofibril structure of D30 cardiomyocytes through anti-cTnI and anti-α-actinin immunostaining, which can visualize sarcomeric length and Z line, respectively. The majority of cardiomyocytes in *SORBS2*-knockdown group presented a round or oval shape instead of polygonal or spindle-like outlines in control group (Figure 3A). Close lookup of myofilaments showed that the sarcomaric structures in cells with abnormal shapes were disrupted (Figure 3A). Overall, the percentage of cardiomyocytes with well-organized sarcomere and normal shape was much lower in *SORBS2*-knockdown group (Figure 3B). In addition, we examined the ultrastructural organization of sarcomere of D30 cardiomyocytes by transmission electron microscopy. The disrupted sarcomeres were also noted in *SORBS2*-knockdown cardiomyocytes (Figure 3C).

**Figure 3.**
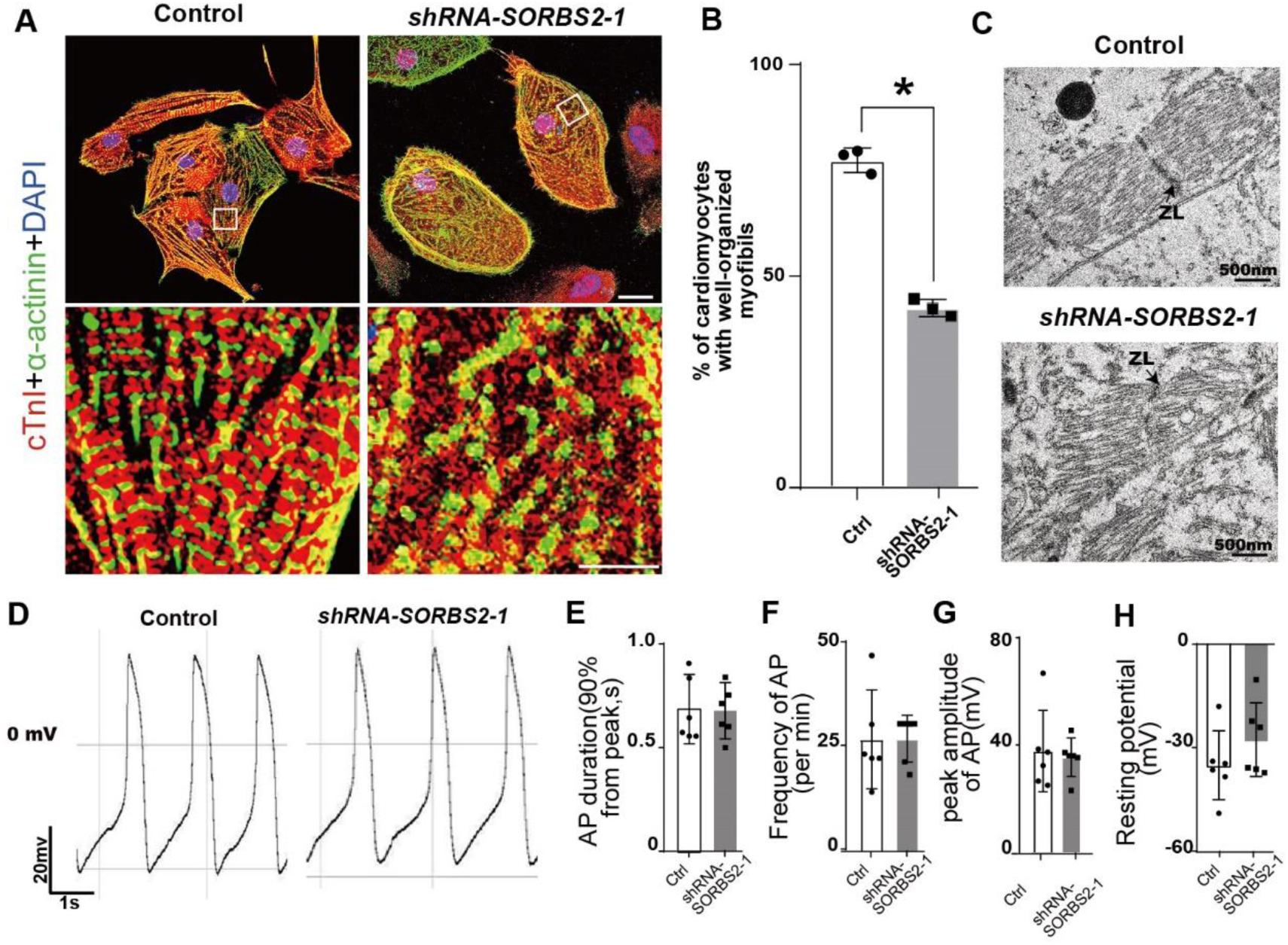
Defective sarcomeric structure but normal electrophysiology of *SORBS2*-knockdown cardiomyocytes. A. Immunostaining of D30 cells with anti-cardiac troponin I (cTnl, red) and anti-α-actinin (green) antibodies. Boxed areas are magnified in the lower panels. Scale bar: 25μm in upper panel and 5μm in lower panel. B. Quantification of cardiomyocytes with well-organized sarcomeres (Control: n=211, *SORBS2-knockdown*: n=197). **, *p* <0.01. C. Transmission electron microscopy images of D30 cardiomyocytes. ZL, Z line. D. Representative action potentials of hESCs-derived ventricular-like cardiomyocytes. E-H. Quantification of action potential parameters of ventricular-like cardiomyocytes (Control, n =6; *SORBS2-knockdown*, n = 6). Frequency (E), action potential duration (F), the peak amplitude (G), and the resting membrane potential (H).

The weakened beating force of *SORBS2*-knockdown cardiomyocytes may also be derived from abnormal electrophysiology. To this end, we examined the electrical activities of dissociated D30 cardiomyocytes by patch clamping. Three types of spontaneous action potentials (ventricular-like, atrial-like, and nodal-like) were all observed in both control and *SORBS2*-knockdown cardiomyocytes. The dominant type of cardiomyocytes is ventricular-like (Figure 3D). Statistical analyses on action potential parameters of ventricular-like cells, including average action potential (AP) duration at 90% repolarization, average AP frequency, peak amplitude, and resting potential showed no difference between two groups (Figure 3E-3H). These data indicate that *SORBS2*-knockdown did not significantly affect cardiomyocyte electrophysiology, albeit its indispensability in sarcomeric organization.

### *SORBS2* promotes cardiomyocyte differentiation through enhancing SHF fate commitment

Cardiomyocyte differentiation efficiency was assessed by flow cytometry analysis of cTnT^+^ cells. The proportion of cTnT^+^ cells was significantly decreased in *SORBS2*-konckdown group at D15 (Figure 4A and 4B). It indicates that *SORBS2* might have an early role in regulating cardiomyocyte differentiation in addition to its role in maintaining sarcomeric integrity of cardiomyocytes. Consistently, we detected significantly decreased expression of sarcomeric genes like *TNNT2, MLC-2A, MYH6* and *MYH7* upon the onset of cardiomyocyte formation in *SORBS2*-konckdown group (Figure 4C-4F).

**Figure 4.**
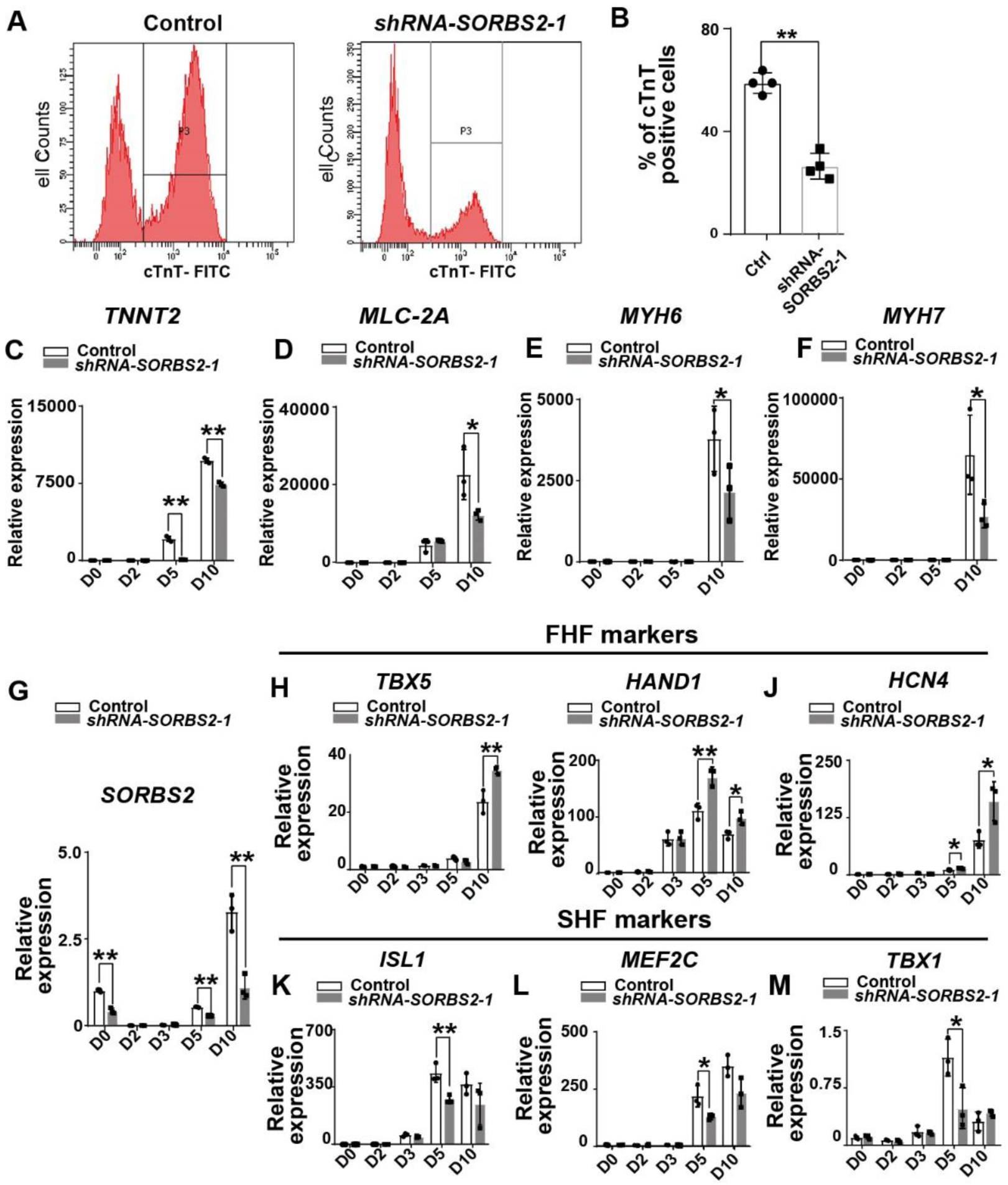
Impair cardiomyocyte differentiation and SHF specification in *in vitro* cardiogenesis of *SORBS2*-knockdown hESCs. A. Flow cytometry analysis of cardiomyocytes at D15. P3 indicates cTnT^+^ population. B. Quantification of cTnT^+^ cells (n = 4). **, *p* <0.01. C-F. qPCR quantification of cardiogenic marker expression at different differentiation time points (n = 3 for each time point). *, *p* <0.05, **, *p* <0.01. G. qPCR quantification of *SORBS2* expression dynamics (n = 3 for each time point). **, *p* <0.01. H-M. qPCR quantification of cardiac progenitor marker expression at different time points (n = 3 for each time point). *, *p* <0.05, **, *p* <0.01.

We examined *SORBS2* expression dynamics to help identify the critical time when *SORBS2* regulates cardiomyocyte differentiation. *SORBS2* was expressed in hESCs, but the expression was quickly turned off at the mesodermal cell stage (D2-D3). Interestingly, *SORBS2* began to re-express at the cardiac progenitor stage D5 and continuously increased with cardiomyocyte differentiation (Figure 4G). In *SORBS2*-knockdown cells, *SORBS2* expression was significantly lower at all the time points (Figure 4G). As previously shown, *SORBS2*-knockdown didn’t significantly affect hESC pluoripotency. In the initial mesodermal commitment stage, the expression levels of mesoderm markers, *T* and *MESP1*, were not changed by *SORBS2*-knockdown (Figure S3A-S3B), which was consistent with absent *SORBS2* expression at this stage.

The reactivation of *SORBS2* expression in cardiac progenitor specification stage indicated a potential role in regulating cardiac progenitor formation. There are two sets of molecularly distinct cardiac progenitors during mammalian heart development, referred to as the first and second heart fields (FHF and SHF). These two populations share common ancestors and contribute to distinct anatomical structures of heart (27, 28). Interestingly, we found significantly increased expression of FHF markers (*TBX5, HCN4, HAND*1) while significantly decreased expression of SHF markers (*TBX1, ISL1, MEF2C*) in *SORBS2*-knockdown cells (Figure 4H-4M). The inverse correlation of FHF and SHF markers suggests that *SORBS2* is a critical regulator to maintain the balance between FHF and SHF cells when derived from common progenitors. SHF gives rise to cardiac outflow tract, right ventricle, and inflow tract (28). Defects in these embryonic structures often lead to CHDs commonly seen in 4q deletion syndrome.

### Transcriptomic analysis reveals signaling pathway alterations that tilt cardiac progenitor specification towards SHF

To understand how *SORBS2* regulates cardiac progenitor differentiation, we collected D5 cells for RNA-seq. There were 1406 differentially expressed genes between two groups (padj < 0.05, Figure 5A). Using a stringent threshold (padj. < 0.05, |fold change| > 2), we selected out 160 down-regulated and 104 up-regulated genes (Table S2-S3) for Go enrichment analysis. Results showed that the up-regulated genes were enriched in biological processes like cell adhesion, and collagen fibril organization etc. (Figure 5B), which might be a compensatory reaction to reduced SORBS2 as a cytoskeleton component. The down-regulated genes were enriched in biological processes like heart development, pattern specification process etc. (Figure 5C). It suggests that *SORBS2* positively regulates cardiac progenitor specification.

**Figure 5.**
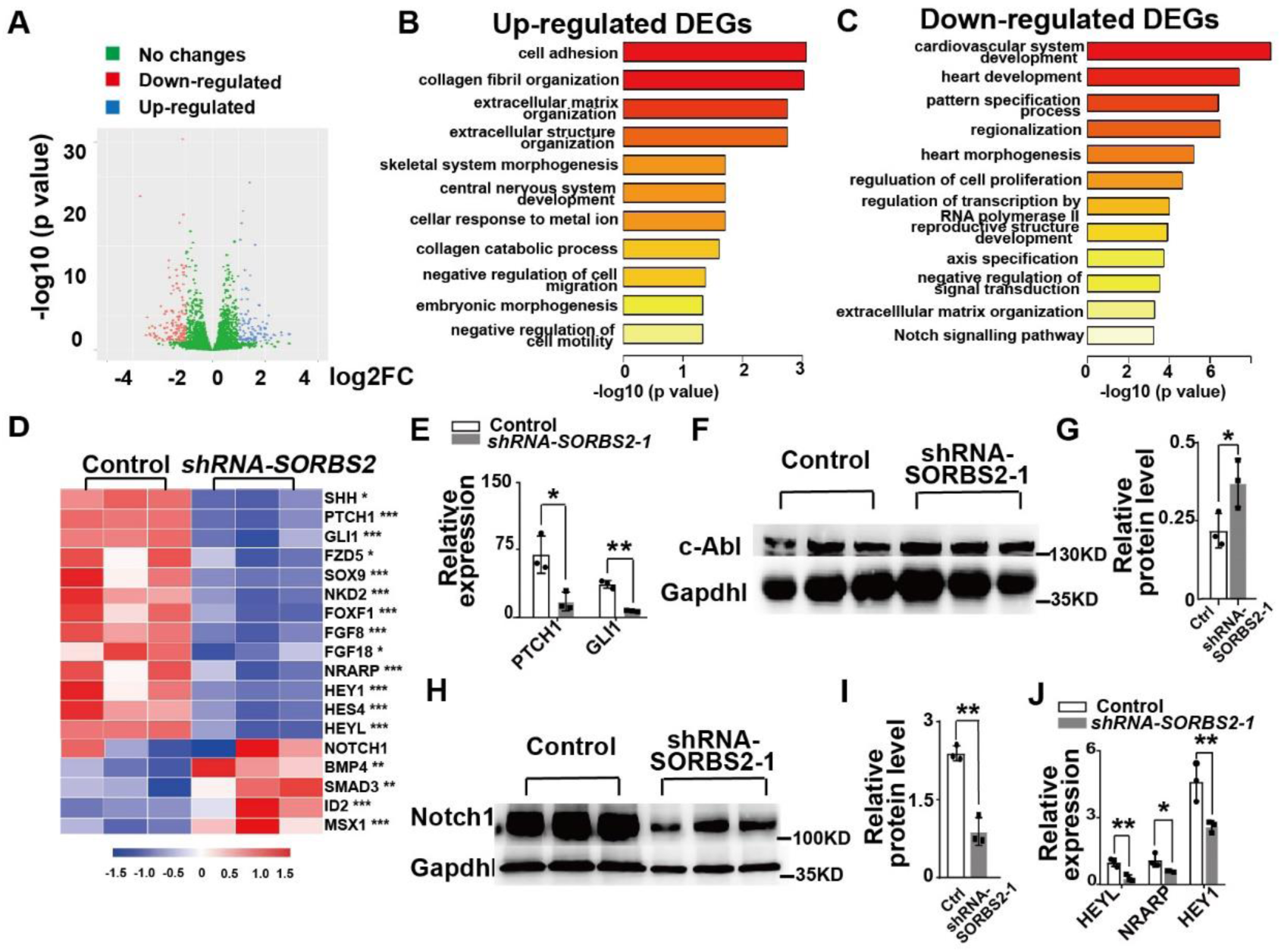
SORBS2 promotes SHF specification through c-ABL/NOTCH/SHH axis. A. Volcano plot illustrates the differential gene expression from D5 RNA-seq data. Orange, down-regulated genes. Blue, up-regulated genes. (|log2(fold change)| > 1 and *padj.* <0.05) B- C. GO biological process analysis of differentially expressed genes. Up-regulated pathways (B). Down-regulated pathways (C). D. Heatmap illustrates gene expression changes of critical signaling pathways. Color tints correspond to expression levels. *, *padj*<0.05. **, *padj*<0.01. ***, *padj*<0.001. E. qPCR quantification of SHH signaling targets at different time points (n = 3 for each time point). *, p <0.05, **, p <0.01. F. Western blot analysis of c-ABL expression in D5 cells. G. Quantification of Western blot (n=3). *, p <0.05. H. qPCR quantification of NOTCH1 signaling targets at different time points (n = 3 for each time point). *, p <0.05. I. Western blot analysis of NOTCH1 expression in D5 cells. J. Quantification of Western blot (n=3). **, p <0.01. K. Diagram depicting the role of *SORBS2* in cardiogenesis.

We showed earlier that *Sorb2*^*-/-*^ mice had ASD. Previous studies have shown that Shh signaling is necessary for the proper differentiation and migration of atrial septal progenitors in posterior SHF to form the primary atrial septum (27, 29). As expected, we noticed the reduced expression of *SHH* and SHH signaling targets *PTCH1* and *GLI1* (Figure 5D). These results were further confirmed by qPCR (Figure 5E). In addition, we also noted reduced expression of genes (*FZD5, SOX9, NKD2, FOXF1*) involved in canonical WNT signaling and elevated expression of genes (*BMP4, SMAD3, ID2, MSX1*) involved in BMP signaling (Figure 5D). Inhibition of canonical WNT signaling and upregulation of BMP signaling promote FHF progenitor formation and meanwhile impair SHF progenitor differentiation(30). Therefore, these expression changes may collectively contribute to the disrupted balance of FHF and SHF progenitors. We confirmed these results by qPCR (Figure S3C). FGF signaling is required for both FHF and SHF progenitor expansion (28, 31). We noted the reduced expression of *FGF8* and *FGF18* in *SORBS2*-knockdown cells (Figure 5D), which may contribute to the reduced efficiency of cardiomyocyte differentiation. We also validated these changes by qPCR (Figure S3C).

### SORBS2 promotes SHH signaling through inhibiting c-ABL-mediated NOTCH1 degradation

To understand how SORBS2 regulates SHH signaling, we looked for SORBS2 function besides its role as a cytoskeleton component. SORBS2 can interact with the non-receptor tyrosine kinase c-ABL as SH3 domain-containing adaptor (21). The binding of SORBS2 to c-ABL triggers the recruitment of ubiquitin ligase CBL and leads to the ubiquitination of c-ABL (32). Indeed, we noted that c-ABL protein level was significantly elevated in *SORBS2*-knockdown cells (Figure 5F-5G). c-Abl can promote Notch endocytosis to modulate Notch protein level. Hence we examined NOTCH1 protein level at D5 cells. As expected, NOTCH1 protein level decreased significantly in *SORBS2*-knockdown group (Figure 5H-5I). Consistently, we noted that the expression of NOTCH signaling target genes *HEY1, HEYL, NARAP* and *HES4*, was also significantly reduced in RNA-seq and GO analysis (Figure 5C-5D). The expression reduction of *HEY1, HEYL* and *NARAP* was further validated by qPCR (Figure 5J). In contrast, there was no change in the transcriptional level of NOTCH1 (Figure 5D), suggesting that the regulation of SORBS2 on NOTCH1 signaling is through modulating protein level. Notch signaling is a well-known molecular mechanism promoting Smo accumulation in cilia and enhancing cellular response to Shh (33, 34). Therefore, the reduced SHH signaling in *SORBS2*-knockdown cells may be a consequence of the decreased NOTCH1 activity.

### Rare *SORBS2* variants are significantly enriched in CHD patients

To validate the genetic association of previously identified CNV genes with CHD, we performed targeted sequencing on 300 complex CHD cases. Besides 23 candidate CNV genes containing *SORBS2*, the targeted panel also includes 81 known CHD genes from literature. For 298 CHD patients included in our study, 98% of the target regions were covered at least 100 times. 220 subjects of Han Chinese descent without CHD from the 1000 genome project were used as controls. The ethnic backgrounds of cases and controls were compared by principal component analysis (PCA) with SNP genotype data from all the participants of this study. Results indicated that CHD cases were clustered together with controls, indicating a matched ethnicity between two groups (Figure S4). A total of 1560 exonic variants from CHD and controls passed quality control and were included for further analyses. 54.29 % of these variants (n=847) were nonsynonymous SNV, 0.38% (n=6) stop-gain and 45.32% (n= 707) synonymous (Figure 6A). Distribution of exonic variants in the breakdown categories of CHD and control groups was shown in Table S4.

**Figure 6.**
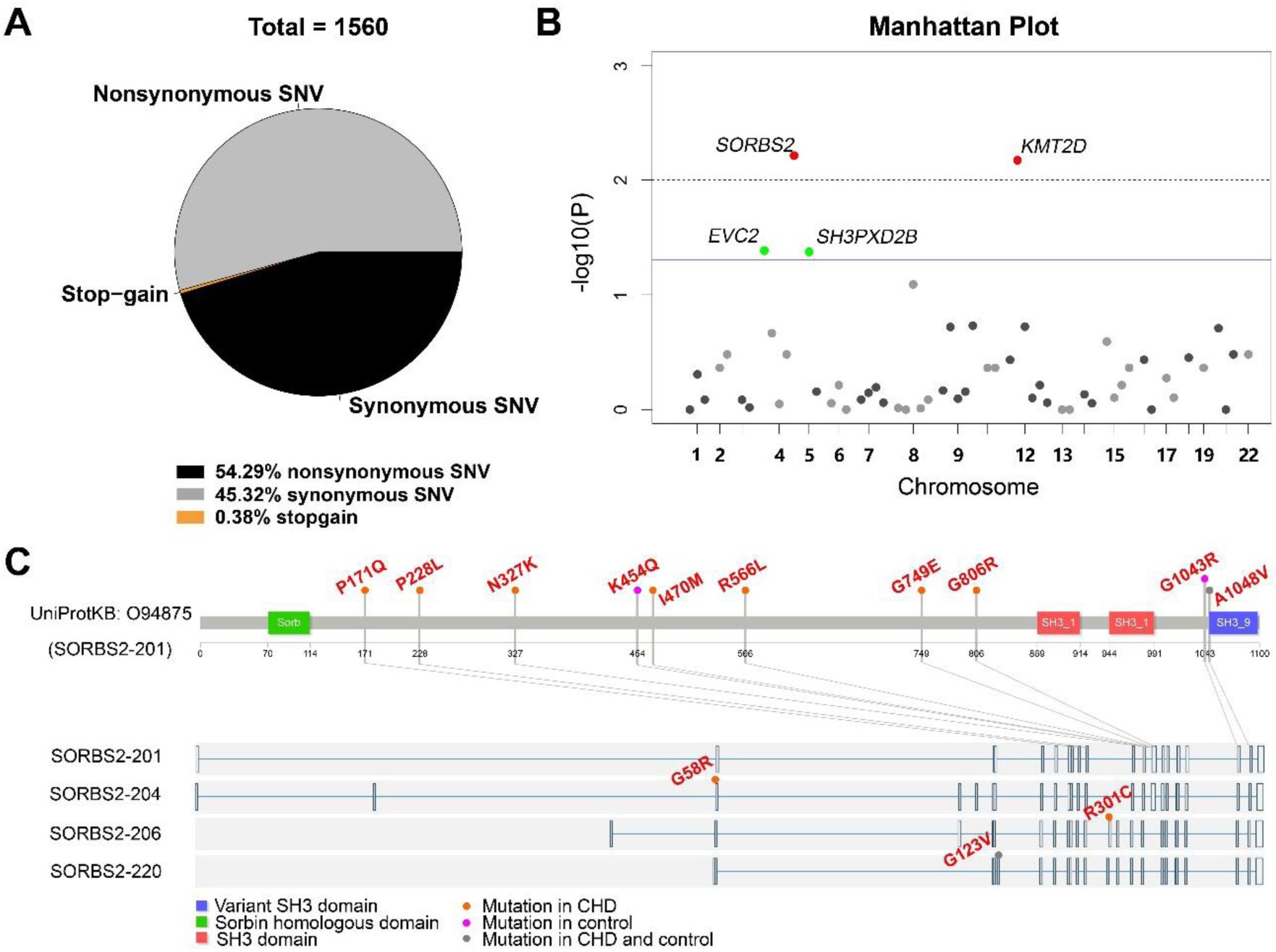
Rare *SORBS2* variants are significantly enriched in CHD patients. A. Descriptive statistics of the identified exonic variants. B. Manhattan plot of gene level Fisher’s exact test of rare damaging variant counts between CHD and control groups. Raw *p* values of 0.05 and 0.01 were indicated by a blue line and a grey dash line, respectively. Genes (*SORBS2, KMT2D*) with *q* value lower than 0.2 were highlighted in red. Genes *(EVC2, SH3PXD2B*) with *p* < 0.05 but q > 0.2 were highlighted in green. C. Illustration of rare damaging variants in *SORBS2*. Most variants are indicated in the main *SORBS2* isoform (SORBS2-201). Three isoform-specific variants are shown in the corresponding exons. Variants in the CHD or control groups are indicated by orange or pink dots, respectively. Variants appearing in both groups are indicated by grey dots.

In current CHD disease model, rare variants contribute significantly to the pathogenesis of CHD (3). Of the 847 nonsynonymous variants, 43.57% (n=369) variants had a minor allele frequency (MAF) below 1% across the public control database (Exome Aggregation Consortium, ExAC), and were adjudicated as “damaging” by at least two algorithms (PolyPhen2, SIFT, or MutationTaster). These rare damaging variants are located in 81 genes, including 65 known CHD genes and 16 candidate genes. We applied gene-based statistic tests to evaluate the cumulative effects of rare damaging variants (MAF<1%) on CHDs. Genes with at least two rare damaging variants (n=57) were included for analysis (Table S5). One-tailed Fisher’s exact test was performed to compare the mutational load of each gene between two groups. 4 out of 57 genes (*SORBS2, KMT2D, EVC2 and SH3PXD2B*) had a *p*-value lower than 0.05 (one-tailed Fisher’s exact) and 2 of them (*SORBS2* and *KMT2D)* had a statistically significant mutation burden after the correction for multiple testing (q<0.20) (Figure 6B, Table S5). *KMT2D* is a well-established CHD gene (35, 36). Our data showed that *SORBS2* variants had similar level of enrichment in CHDs as the known CHD genes. The distribution of rare *SORBS2* damaging variants in CHD patients spread throughout the gene (Figure 6C). Although we didn’t detect *SORBS2* nonsense variants in CHD patients, the excessive burden of *SORBS2* missense mutations in CHD group suggests a role in CHD development. Indeed, the majority of patients with rare *SORBS2* variants had ASD (Table S6), the defect seen in *Sorbs2*^*-/-*^ mouse heart. Overall, these data further support that *SORBS2* contributes to the pathogenesis of cardiac defects of 4q deletion syndrome.

## Discussion

Chromosome 4q deletion syndrome is a genetic disease resulting from chromosomal aberration that causes a portion of chromosome 4 long arm missing (20). Chromosomal deletions are interstitial or terminal. Although deletion size and position are highly variable among patients, some breakpoints occur more frequently. With the advent of high-resolution chromosomal array, more submicroscopic microdeletions are being identified, particularly the clustering terminal 4q deletions(19, 37). These microdeletions greatly facilitate the identification of disease genes underpinning various defects. 4q deletion syndrome patients have a spectrum of clinical manifestations including craniofacial, skeletal, cardiovascular, gastrointestinal abnormalities, mental and growth deficiencies (20). Among them, CHD is a common defect seen in about half patients (20). A study narrowed the cardiovascular critical region to 4q32.2–q34.3, which contains *TLL1, HPGD* and *HAND2* that participate in cardiovascular development (38). Over-represented right-sided CHDs in 4q deletion syndrome patients suggest that *HAND2*, an essential regulator of SHF development, is mainly responsible for CHD phenotype (38, 39). However, a portion of 4q deletions associated with CHD that we and others have discovered do not cover this defined region(19, 37, 40, 41). *SORBS2* within terminal 4q deletions has been proposed as a candidate CHD gene due to a rare small deletion event and its high expression level in heart (37). The follow-up work to sequence CHD patients for *SORBS2* mutations, stated in this article, has never been published. It leaves this single-case association in an uncertain situation. Here we presented evidence from animal model, in vitro cardiogenesis and mutation analysis to demonstrate that SORBS2 functions not only as a sarcomeric component to maintain cardiomyocyte function, but also as an adaptor protein to promote SHF development through c-ABL/NOTCH/SHH axis and its haploinsufficiency may contribute to the overall pathogenicity of chromosome 4q deletion.

The common CHDs in 4q deletion syndrome include ASD, ventricular septal defect (VSD), pulmonary stenosis/atresia, and tetralogy of Fallot etc. (20, 42, 43). The affected structures are atrial septum and cardiac outflow tract, which are derived from posterior and anterior SHF, respectively (27, 44). Although ASD is detected in *Sorbs2*^*-/-*^ mouse hearts, which supports that *Sorbs2* regulates SHF development *in vivo* and the role is conserved across species, no conotruncal defect is present in mouse mutant hearts. It is different from the high penetrance of conotruncal defects seen in human patients. The discrepancy may be due to genetic modifiers that, together with *SORBS2* haploinsufficiency, cause the developmental defect in cardiac outflow tract. An obvious genetic modifier is the *HAND2* gene, which is co-missing with *SORBS2* in large 4q deletions. Previous studies have shown that CHD is observed more frequently in patients with the terminal deletion at 4q31, which includes *HAND2*, than in patients with the terminal deletion at 4q34 or 4q35 (42, 43). *Hand2* is essential for anterior SHF progenitor to proliferate and survive (45) and functions downstream of Notch signaling to regulate cardiac OFT development (46). Therefore, *SORBS2* and *HAND2* may have synergic effects at both SHF development and NOTCH signaling levels to control OFT development. *HELT* within 4q35.1 might be another genetic modifier of *SORBS2*, particular for deletion not including *HAND2. Helt* encodes a Hey-related bHLH transcription factor that is expressed in both brain and heart, and mediates Notch signaling (47). Therefore, *SORBS2* and *HELT* might synergistically regulate OFT development through modulating NOTCH signaling. The relatively high incidence of rare *SORBS2* variants in CHD group suggests that it may act as a genetic contributor to the pathogenesis of isolated CHDs, albeit a low penetrance by itself. Our findings show that *SORBS2* regulates SHH and NOTCH signaling. It might interact with other components of SHH and NOTCH signaling pathways to modify cardiac phenotype, which is consistent with a recent report indicating cilia-related genes are significantly enriched for damaging rare recessive and compound heterozygous genotypes(48).

There are two types of ASD in *Sorbs2*^*-/-*^ heart, including atrial septal aplasia/hypoplasia and DAS. It is surprising that the opposite cellular changes occur in the same type of mutants. Lineage tracing has shown that primary atrial septum is mainly derived from Shh-responsive posterior SHF progenitors (27, 29). Knockout of Shh receptor *Smo* in SHF or Shh-responsive cells impairs the differentiation and migration of SHF progenitors and consequently leads to atrial septal aplasia (27, 29). In *Smo* mutants, more Shh-responsive progenitor-derived cells are directed into atrial free wall instead of primary atrial septum(29). *Sorbs2*-knockdown tunes down Notch signaling. It is well established that Notch signaling facilitates Smo accumulation in primary cilia (33, 34). Hence *Sorbs2* knockout may dampen Shh signaling through reducing Smo level within primary cilia. In comparison to *Smo* mutant mice, the extent of Shh signaling reduction in *Sorbs2*^*-/-*^ mice may be much milder, which is supported by the phenotypic difference between two mutants. There is severe cardiac OFT defect in *Smo* mutants but not in *Sorbs2*^*-/-*^ mice. If the migration of Shh-responsive SHF progenitor is more dependent on Shh signaling than progenitor differentiation does, Shh signaling dampening would hit cell migration first before it can affect their differentiation. In such a condition, SHF progenitors in *Sorbs2*^*-/-*^ mice might have normal cell number but abnormal cell migration. If the direction of unguided migration is randomized, there is a small chance that more Shh-responsive SHF progenitors, including those destined for atrial free wall, migrate into atrial septal area, thereby producing an extra atrial septum in a small percentage of *Sorbs2*^*-/-*^ embryos.

DAS, also called Cor triatriatum type C in the original report (49), is a very rare CHD characterized by an extra septal structure to the right side of primary atrial septum (50). The anatomic abnormality is implicated as a cause of paradoxical thromboembolic event to stroke or heart attack (51, 52). However, its etiology and pathogenesis are entirely unknown. Our data reveal the first genetic etiology of DAS. Interestingly, Cor triatriatum, another type of abnormal atrial septation, have been reported in a patient with a terminal 4q34.3 deletion (53), which includes *SORBS2* but not *HAND2*. It has been speculated that the additional atrial septum might result from persistence of embryologic structures or abnormal duplication of atrial septum. The impaired cardiogenesis in *SORBS2*-knockdown cells suggests that the latter scenario may be the underlying pathogenesis.

Cardiomyocytes are derived from two different cardiac progenitor populations, FHF and SHF. These two populations are derived from Mesp1^+^ mesodermal progenitors during gastrulation (54, 55). In *in vitro* cardiogenesis, molecularly distinctive FHF and SHF progenitors simultaneously appear during differentiation process (56). However, what controls the fate commitment to FHF or SHF remains unclear. *SORBS2* is not expressed at mesodermal stage, but its expression is switched on at cardiac progenitor stage. The temporal expression pattern suggests a role in the fate choice of cardiac progenitor. Interestingly, *SORBS2*-knockdown causes a tilted balance between FHF and SHF progenitors, which leans towards FHF fate. It is similar to a molecular switch in the chordate model Ciona. Tbx1/10-Dach pathway maintains the separate lineages of FHF and SHF in Ciona. Deletion of *Tbx1/10* or *Dach* causes ectopic FHF marker expression in SHF cells (57). Interestingly, a recent study has shown that *SORBS2* mutation is associated with arrhythmogenic right ventricular cardiomyopathy (58), which is considered as a disease of disrupted differentiation of cardiac progenitor cells(59).

Overall, our data reveal that *SORBS2* has a dual role in regulating cardiac development and function. The discovered role of *SORBS2* in promoting SHF development substantiates that *SORBS2* haploinsufficiency contributes to CHD seen in 4q deletion syndrome.

## Material and Methods

### Mouse lines and breeding

Mice were housed under specific pathogen-free conditions at the animal facility of Shanghai Children’s Medical Center. *Sorbs2*^*flox/flox*^ mice (24) were a gift from Dr. Guoping Feng’s lab (McGovern Institute for Brain Research, MIT, Cambridge). *Sorbs2*^*-*^ allele was obtained by breeding *Sorbs2*^*flox*^ allele into CMV-Cre mouse (60). The strains were backcrossed with C57BL/6 to maintain the lines ever since we obtained them. 2-6 months old males and females were used for timed mating and embryos were collected at E18.5. Neither anesthetic nor analgesic agent was applied. CO2 gas in a closed chamber was used for euthanasia of pregnant dam and cervical dislocation was followed. Isolated fetuses were euthanized by cervical dislocation. Animal care and use were in accordance with the NIH guidelines for the Care and Use of Laboratory Animals and approved by the Institutional Animal Care and Use Committee of Shanghai Children’s Medical Center.

### Histological analysis

For cardiac phenotype analysis, embryos were collected at E18.5 and were fixed in 10% formalin overnight. Isolated hearts were processed for paraffin embedding, sectioned at a thickness of 4 μm and stained with Hematoxylin and Eosin. Stained sections were imaged using a Leica DM6000 microscope.

### H1 hESCs cell cultures and cardiomyocytes differentiation

Undifferentiated H1 hESC lines were maintained in a feeder-free culture system. Briefly, we precoated the well plates with Matrigel (354277, BD Biosciences), and then seeded and cultured cells with TeSR(tm)-E8(tm) medium (05840,Stemcell). When cells reached 80% confluence, they were passaged routinely with Accutase (07920, Stemcell). For cardiomyocyte differentiation, cells were induced using a chemically defined medium consisting of three components (CDM3): the basal medium RPMI 1640 (C14065500, Gibco), L-ascorbic acid 2-phosphate(213µg/ml,113170-55-1,Sigma) and Oryza sativa-derived recombinant human albumin(500µg/m,HY100M1,Healthgen Biotechnology Corp). In brief, single cell suspensions were prepared using Accutase and were seeded in 12-well Matrigel-coated plate at a density of 4×10^5^ cells/well. When cells reached 80%-90% confluence (Day 0), cells were fed by 2ml CDM3 basal medium supplemented with CHIR99021 (6μM, 72052, Stem cell). 48 hours later (Day 2), medium was replaced with 2ml CDM3 supplemented with Wnt-C59 (2μM, 1248913, Peprotech Biogems). After 96 hours (day 4), the medium was replaced with CDM3 basal medium every other day until the appearance of cell beating.

### Lentiviral shRNA knockdown of *SORBS2* in hESCs

shRNA plasmid vectors (U6-MCS-Ubiquitin-Cherry-IRES-puromycin) expressing target-specific sequences against human *SORBS2* and non-target scrambled control were purchased from Shanghai Genechem Co. The targeting sequences of *SORBS2* shRNA-1 and shRNA-2 are 5′-TCCTTGTATCAGTCCTCTA-3′ and 5′-TCGATTCCACAGACACATA-3′, respectively. The sequence of scrambled shRNA is 5′-TTCTCCGAACGTGTCACGT-3′. Lentiviral particles were produced by transfecting human embryonic kidney (HEK) 293FT cells with shRNA, psPAX2 (#12260, Addgene), and pMD2.G (#12259, Addgene) plasmids. Efficiency of gene knockdown was examined using qRT-PCR.

### Western blot

Cells were lysed in RIPA buffer (P0013B, Beyotime) containing protease inhibitors (P1010, Beyotime). Protein concentrations were determined with the BCA™ protein assay kit (Thermo). Protein was separated via 8% SDS-PAGE and afterwards transferred to polyvinylidene difluoride (PVDF) membrane. After blocking by 5% non-fat milk for 1 hour, primary antibodies were incubated overnight at 4°C. The membrane was washed with TBST and incubated with second antibodies for 0.5 hour at room temperature. Bands were detected with the Immobilon ECL Ultra Western HRP Substrate (WBULS0500, Sigma) and band intensity was analyzed by ImageJ software. Antibodies: c-ABL (1:1000; A0282, Abclonal), NOTCH1 (1:1000; 3608s, CST), GAPDH (1:1000; ab8245, Abcam). Goat Anti-Rabbit IgG H&L (HRP) (1:5000; ab6721, Abcam), Goat Anti-Mouse IgG H&L (HRP) (1:5000; ab205719, Abcam).

### Immunofluorescent staining

Cells were fixed in 4% paraformaldehyde for 10 min, permeabilized in 0.5% Triton X-100/PBS for 20 min, and then blocked in 5% bovine serum albumin/PBS for 30 min. Fixed cells were stained with the following primary antibodies: TRI-1-60 (1:200; Ab-16288, Abcam), OCT4 (1:200; Ab-18976, Abcam), SOX2 (1:200; Ab-80520, Abcam), cTnI (1:200; Ab-92408, Abcam) and α-actinin (1:200; A-9776, Sigma). These primary antibodies were visualized with AlexaFluor 488 (1:1000; A-11029, Invitrogen) or AlexaFluor 633 (1:1000; A-21071, Invitrogen). Nuclei were stained with DAPI. Fluorescence images were acquired using a Laser confocal microscope (Leica TCS SP8).

### Flow cytometric analysis

In brief, D15 cells were harvested in 0.25% trypsin/EDTA at 37 °C for 15mins and subsequently neutralized by 10% FBS in DMEM. Then cells were centrifuged at 1000 rpm for 5min and resuspended in Invitrogen™ FIX&PERM solution and kept at 4°C for 30min. After washing, cells were incubated with anti-cTnT antibody (1:100; ab105439, Abcam) in washing buffer on ice in the dark for 45 min. Cells were centrifuged, washed and resuspended for detection. Data were collected by BD FACSCanto™ flow cytometer and analyzed by BD FACS software.

### Electron Microscopy (EM)

D30 cardiomyocytes were harvested using 0.25% trypsin/EDTA and prefixed with 2.5% glutaraldehyde in 0.2M phosphate buffer overnight at 4°C. Samples were washed and then post-fixed with 1% osmium tetroxide for 1.5 hours. Next, cells were routinely dehydrated in an ethanol series of 30%, 50%, 70%, 80% and 95% ethanol for 15 min each, 100% ethanol and acetone twice for 20 min each at room temperature and then embedded in epoxy resin. Sections (70 nm thick) were poststained in uranyl acetate and lead citrate and visualized on Hitachi 7650 microscope.

### Electrophysiological recordings

D30 h1ESCs-CMs were digested by Accutase (7920, STEMCELL Technologies, Canada), washed cells one time with baseline extracellular fluid and then moved to the stage of an inverted microscope (ECLIPSE Ti-U, Nikon, Japan) for patch-clamp recording. h1ESCs-CMs were continuously perfused by extracellular solution through a “Y-tube” system with solution exchange time of 1 minute. Whole-cell patch-clamp recordings were performed using Axopatch 700B (Axon Instruments, Inc., Union City, CA, USA) amplifiers under an invert microscope at room temperature (22-25°C). Glass pipettes were prepared using borosilicate glasses with a filament (Sutter Instruments Co, Novato, CA, USA) using the Flaming/Brown micropipette puller P97 (Sutter Instruments Co, Novato, CA, USA). The final resistance parameters of patch pipette tips were about 2-4 MΩ after heat polish and internal solution filling. After forming “gigaseal” between the patch pipette and cell membranes, a gentle suction was operated to rupture the cell membrane and establish whole-cell configuration. All current signals were digitized with a sampling rate of 10 kHz and filtered at a cutoff frequency of 2 kHz (Digidata 1550A, Axon Instruments, Inc., Union City, CA, USA). The spontaneous action potentials were recorded by gap free mode with a sampling rate of 1 kHz and filtered at a cutoff frequency of 0.5 kHz. If the series resistance was more than 10 MΩ or changed significantly during the experiments, the recordings were discarded from further analysis. The pipette internal solution for action potential recording contained (in mM): KCl 150, NaCl 5, CaCl2 2, EGTA 5, HEPES 10, MgATP 5 (pH 7.2, KOH), and baseline extracellular solution and extracellular solution for action potential recording contained (in mM): NaCl 140, KCl 5, CaCl2 1, MgCl2 1, Glucose 10, HEPES 10 (pH 7.4, NaOH). Data were analyzed by using Clampfit 10.5 and Origin 8.0 (OriginLab, Northampton, MA, USA).

### Quantitative real-time PCR analysis

Total RNA was extracted from D0, D2, D3, D5 and D10 cells using Trizol reagent (Thermo Fisher Scientific, 15596018). Reverse transcription was accomplished with Reverse Transcription Kit (Takara, RR037A) according to the manufacturer’s instructions. qPCR was performed with the SYBR Fast qPCR Mix (Takara, RR430A) in the Applied Biosystems 7900 Real-Time PCR System. Primers are listed in Table S7.

### RNA-Seq

Total RNA of D5 cells was isolated using TRizol reagent (Thermo Fisher Scientific, 15596018). Library preparation and transcriptome sequencing on an Illumina HiSeq platform were performed by Novogene Bioinformatics Technology Co., Ltd. to generate 100-bp paired-end reads. HTSeq v0.6.0 was used to count the read numbers mapped to each gene, and fragments per kilobase of transcript per million fragments mapped (FPKM) of each gene were calculated. We used FastQC to control the quality of transcriptome sequencing data. Next, we compared the sequencing data to the human reference genome (hg19) by STAR. The expression level of each gene under different treatment conditions is obtained by HTSeq-count after standardization. The differentially expressed genes were analyzed by DESeq2 software according to the standard that the *p*-value was under 0.05 after correction and the fold change was more than 2 times. Functional enrichment of differentially expressed genes is analyzed on Toppgene website.

### Network and Gene ontology analysis

From ENCODE database (23), 316 human fetal heart specific genes including *SORBS2* were selected and their expression coefficients were computed. The highly co-regulated transcriptional networks (correlation efficient ≥ 0.8) were constructed and visualized with BioLayout Express3D. The interconnected gene clusters were detected using the MCL (Markov Cluster) algorithm and illustrated with different colors.

### Patient samples

A total of 300 children with complex CHD, including ASD, conotruncal defect etc., were enrolled in our study from November 2011 to January 2014 in Shanghai Children’s Medical Center (Table S8). Patients carrying 22q11.2 deletion and gross chromosomal aberrations were excluded from our study. The mean age of included probands was 10 months with a range of 3 days to 17 years. 188 (62.7%) were boys and 112 (37.3%) were girls. CHD diagnosis was confirmed by reviewing patient history, physical examinations, and medical records. Patients carrying 22q11.2 deletion and gross chromosomal aberrations (e.g., trisomy 21, trisomy 13 and trisomy 18) were excluded from our study. The study conformed to the principles outlined in the Declaration of Helsinki and approval for human subject research was obtained from the Institutional Review Board of Shanghai Children’s Medical Center. Written informed consents were obtained from parents or legal guardians of all patients.

### Control cohort

Exome sequences for 220 control subjects of Han Chinese descent were derived from the 1000 genome project (http://www.internationalgenome.org/data). Raw sequence data in the form of bam or fastq files was re-aligned and re-analyzed with the same bioinformatic pipelines with CHD patients. The ethnicity of cases and controls were investigated by performing PCA with SNP genotype data from all the participants of this study.

### Gene selection and targeted sequencing

We used a customized capture panel of 104 targeted genes, which included 81 known CHD genes (Table S9) and 23 CHD candidate genes (Table S10). The CHD genes were selected through a comprehensive literature search and had been reported by other research groups to be associated with CHD in either human patients or mouse models (1, 61, 62). CHD candidate genes were prioritized from pathogenic CNVs and likely pathogenic CNVs identified in CHD patients in our previous study(19). The coding regions of selected genes and their flanking sequences were covered by the Agilent SureSelect Capture panel and sequenced on the Illumina Genome Analyzer IIx platform according to the protocols recommended by the manufacturers.

### Variant calling and quality control

We used the best practice pipeline of Broad Institute’s genome analysis toolkit (GATK 3.7) to obtain genetic variants from the target sequencing data of 300 CHD cases together with the raw data of 220 Han Chinese control samples. Briefly, Burrows-Wheeler Aligner (BWA, version 0.7.17) was used to align the Fastq format sequences to human genome reference (hg38). De-duplication was performed using Picard and the base quality score recalibration (BQSR) was performed to generate analysis-ready reads. HaplotypeCaller implemented in GATK was used for variant calling in GVCF mode. All samples were then genotyped jointly.

We then excluded the variants based on the following rules: (1) >2 alternative alleles; (2) low genotype call rate <90%; (3) deviation from Hardy-Weinberg Equilibrium in control samples (p < 10^−7^); (4) differential missingness between cases and controls (p < 10^−6^). After alignment and variant calling, we removed 2 subjects with low-quality data, leaving a total of 298 cases and 220 controls.

### Variant enrichment analysis

Given that severe mutations are generally present at low frequencies in the population, we set the relevant variants filtering criteria as follows: (1) variants that located in exonic region, (2) exclude synonymous variants, (3) variants with minor allele frequency (MAF) below 1% according to the public control database (Exome Aggregation Consortium, ExAC), and (4) damaging missense variants predicted to be deleterious by at least two algorithms (Polyphen2 ≥ 0.95/MutationTaster_pred:D/SIFT≤ 0.05). All relevant variants following these criteria were hereafter called “rare damaging” variants. The number of rare damaging variant carriers in each gene was counted in CHD patients and controls. We hypothesized that rare damaging variants should be enriched in CHD patients, hence the carrier and non-carrier groups were compared between CHD patients and controls using a one-tailed Fisher’s exact test. The odds ratio (OR) was calculated. Only genes with at least two variants were retained and multiple testing correction was performed using Benjamini-Hochberg procedure (q-value, adjusted *p*-value after Benjamini-Hochberg testing).

### Statistical analysis

Statistical significance was performed using a two-tailed Student’s t test or χ^2^ test as appropriate. Statistical significance is indicated by *, where *p* < 0.05, **, where *p* < 0.01.

## Acknowledgements

We thank Dr. Guoping Feng’s at Massachusetts Institute of Technology for *Sorbs2*^*flox/flo*x^ mice, Dr. Bingshan Li at Vanderbilt University and Dr. Hao Mei at University of Mississippi Medical Center for advice on genetic data analyses.

## Author contributions

F. L. performed mouse and in vitro cardiogenesis experiments; X. Z., B. W., J. G. and G. Y. did targeted sequencing and analyzed genomic data; H. S. analyzed RNA-seq data; H. S. analyzed CHD phenotype and collected samples. Q. F. and Z.Z. conceived and directed the project. F. L., X. Z. and Z.Z wrote the manuscript. Q. F., H. C. and J. F. commented on the written manuscript.

## Data and code availability

Targeted sequencing raw data of CHD patients have been deposited NCBI’s Sequence Read Archive (PRJNA579193). RNA-seq data have been deposited in NCBI’s Gene Expression Omnibus (GSE137090). R scripts will be available upon request.

