## Supplemental materials for "*SORBS2* is a genetic factor contributing to cardiac malformation of 4q deletion syndrome"

**This file includes:**

**Supplemental Figures**, p2-p5

**Supplemental Videos**, p6

**Supplemental Tables**, p7-p21

### Supplemental Figures

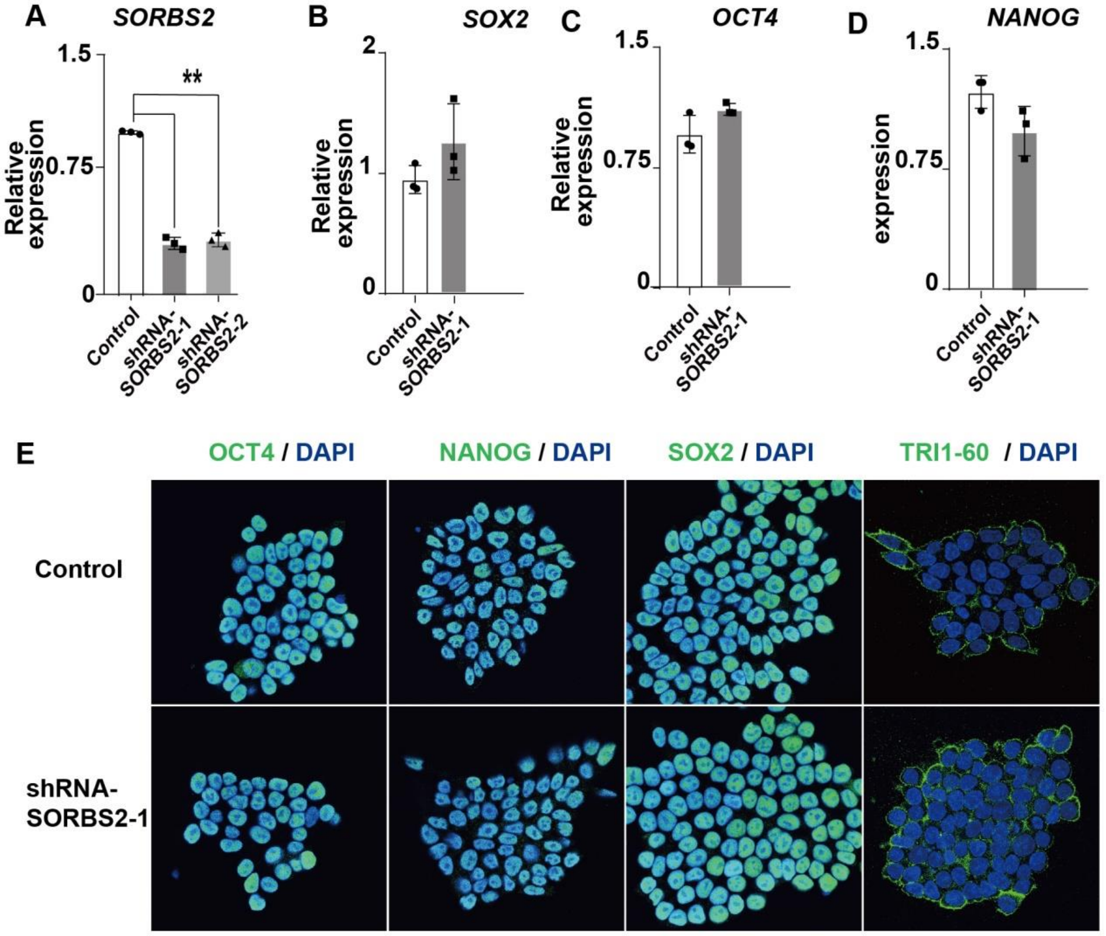


**Figure S1. Characterization of *shRNA-SORBS2* hESCs.**

A. Quantification of *SORBS2* expression level in *shRNA-SORBS2* hESCs (n = 3). **, $p$ <0.01, two-tailed Student's $t$ test.

B-D. Quantification of pluripotential markers expression (n = 3). Two-tailed Student's $t$ test.

E. Immunostaining of hESCs cells with anti-NANOG, anti-OCT4, anti-SOX2 and anti-TRI1-60 antibodies (green).

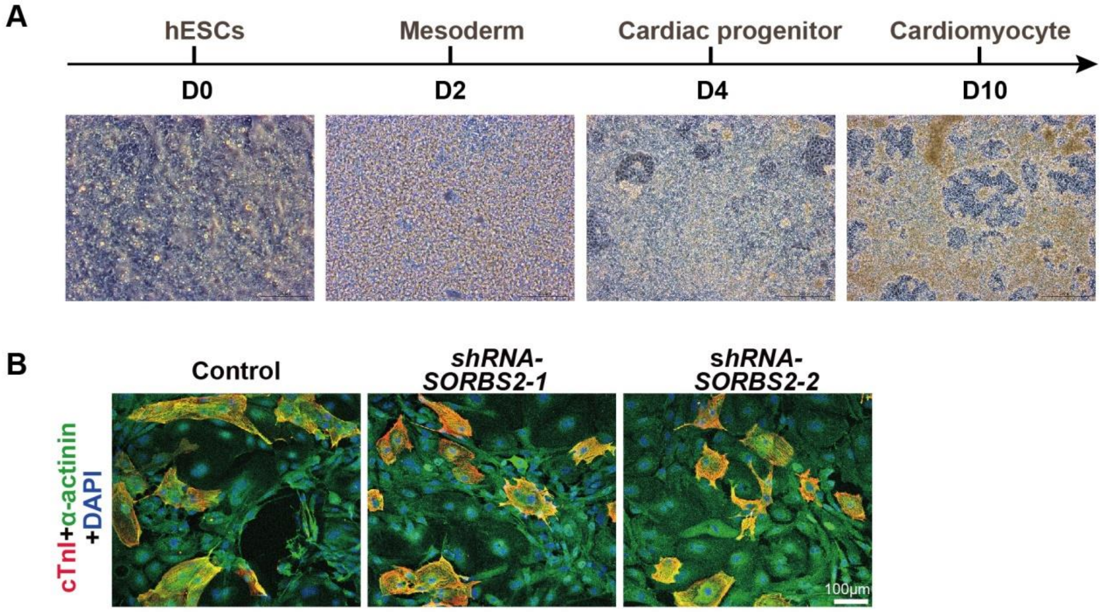


**Figure S2. In vitro cardiogenesis from hESCs.**

A. Cell morphological transition during cardiomyocyte differentiation.

B. Immunostaining of D30 cells with anti-cardiac troponin I (cTnl, red) and anti-α-actinin (green) antibodies.

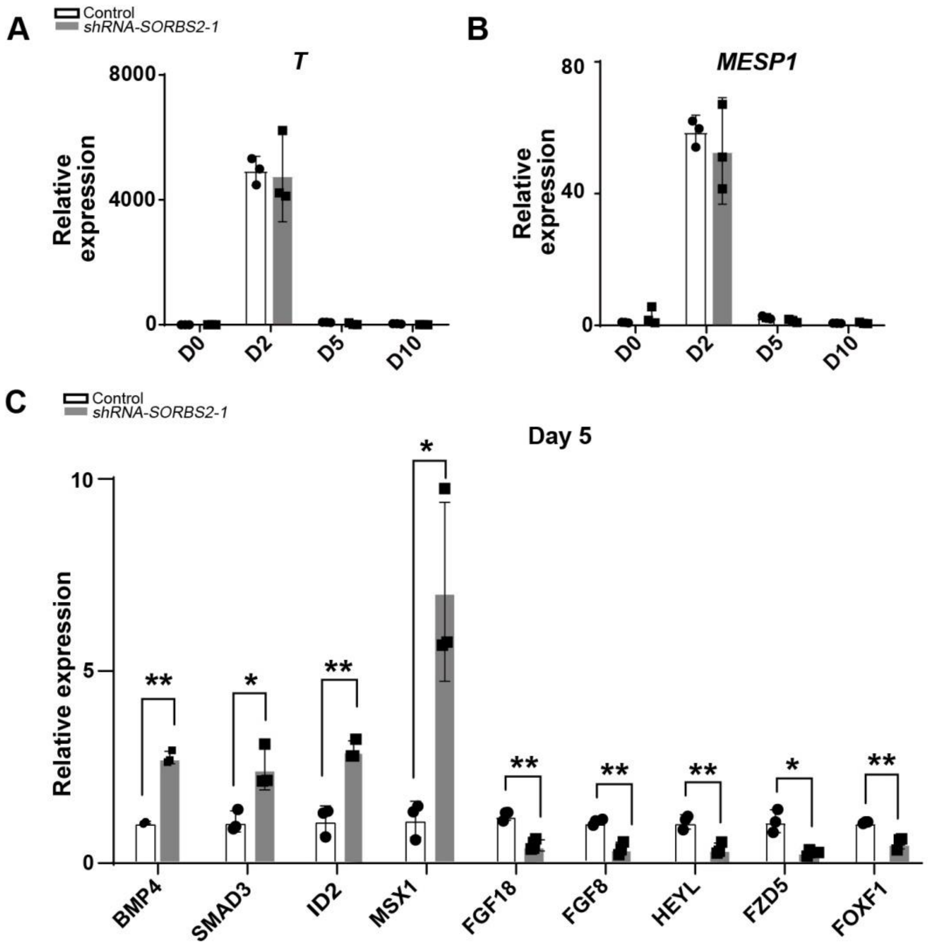


**Figure S3. Molecular profiling of *SORBS2*-knockdown hESCs-derived mesoderm and cardiac progenitors.**

A-B. qPCR quantification of mesoderm markers expression at different differentiation time points (n = 3). Two-tailed Student's t test.

C. qPCR verification of differentially expressed genes in RNA-seq (n=3). *, $p < 0.05$. **, $p < 0.01$. Two-tailed Student's t test.

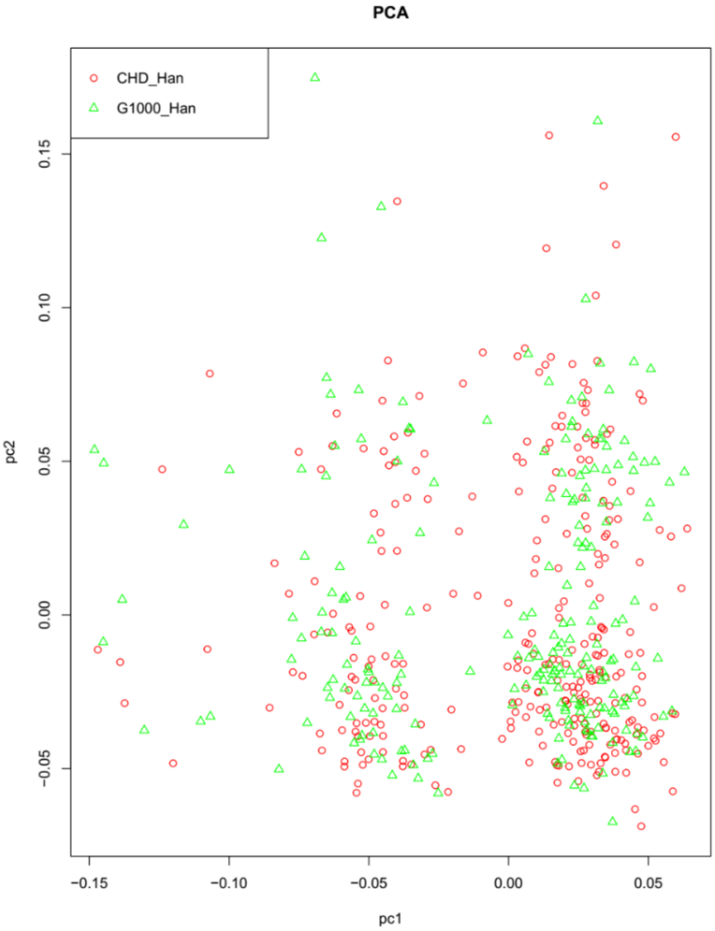


**Figure S4. Ethnic background comparison of CHD and control groups.**

PCA plot of population structure with the top two principle components (PC1: Principle component 1; PC2: Principle component 2). CHD and control samples were clustered together in PCA of SNP genotype.

### Supplemental videos

**Video S1-S3: Videos of beating D20 cardiomyocytes**

S1: control, S2: *shRNA-SORBS2*-1, S3: *shRNA-SORBS2*-2.

### Supplemental tables

**Table S1. Genotyping distribution in embryos from *Sorbs2*$^{+/-}$ mouse intercross**

| Embryonic stage | Total | Genotype of embryos: WT | *Sorbs2*$^{+/-}$ | *Sorbs2*$^{-/-}$ | ASD |
|---|---|---|---|---|---|
| E18.5 | 137 | 42 | 65 | 30[a] | 12[b] |

a: The observed ratio is not different from the expected. Two-sided $\chi 2$ test ($\chi 2$=1.212, *p*=0.546).

b: 10 cases of primary septum hypoplasia/aplasia and 2 cases of double atrial septum.

**The following two tables are provided as independent Excel files:**

**Table S2. Down-regulated genes in *SORBS2*-knockdown D5 cells.**

**Table S3. Up-regulated genes in *SORBS2*-knockdown D5 cells.**

**Table S4. Number of exonic variants detected in CHD and normal controls.**

| | CHD cases | Normal controls |
|---|---|---|
| **All genes** | | |
| Total | 1108 | 842 |
| Synonymous SNV | 506 | 430 |
| Stopgains | 4 | 2 |
| Nonsynonymous SNV | 598 | 410 |
| **CHD genes** | | |
| Total | 899 | 658 |
| Synonymous SNV | 409 | 329 |
| Stopgains | 3 | 2 |
| Nonsynonymous SNV | 487 | 327 |
| **Candidate genes** | | |
| Total | 209 | 184 |
| Synonymous SNV | 97 | 101 |
| Stopgains | 1 | 0 |
| Nonsynonymous SNV | 111 | 83 |

**Table S5. Carriers of rare damaging variants in CHD and normal controls. P.values are calculated with one-tailed Fisher's exact test and q.values are adjusted p-values after Benjamini-Hochberg testing.**

| | Gene | No.carriers in CHD patients (n=298) | No.carriers in controls (n=220) | P.value | Odds Ratio | BH q.value |
|---|---|---|---|---|---|---|
| 1 | *SORBS2* | 20 | 4 | 0.006129 | 3.876383 | 0.1920615 |
| 2 | *KMT2D* | 22 | 5 | 0.006739 | 3.420478 | 0.1920615 |
| 3 | *EVC2* | 13 | 3 | 0.04131 | 3.292905 | 0.6029175 |
| 4 | *SH3PXD2B* | 15 | 4 | 0.04231 | 2.857021 | 0.6029175 |
| 5 | *CHD7* | 7 | 1 | 0.08121 | 5.255293 | 0.925794 |
| 6 | *EHMT1* | 7 | 2 | 0.1859 | 2.617671 | 1 |
| 7 | *PTPN11* | 3 | 0 | 0.1896 | Inf | 1 |
| 8 | *ROR2* | 17 | 8 | 0.1907 | 1.601803 | 1 |
| 9 | *JAG1* | 5 | 1 | 0.1953 | 3.729429 | 1 |
| 10 | *EVC* | 10 | 4 | 0.2162 | 1.872866 | 1 |
| 11 | *FBN1* | 11 | 5 | 0.2558 | 1.646571 | 1 |
| 12 | *ACVR1* | 2 | 0 | 0.3305 | Inf | 1 |
| 13 | *PDLIM3* | 2 | 0 | 0.3305 | Inf | 1 |
| 14 | *SALL4* | 2 | 0 | 0.3305 | Inf | 1 |
| 15 | *TBX1* | 2 | 0 | 0.3305 | Inf | 1 |
| 16 | *NFATC1* | 17 | 10 | 0.3527 | 1.269886 | 1 |

| | | | | | | |
|---|---|---|---|---|---|---|
| 17 | *COL2A1* | 8 | 4 | 0.368 | 1.488552 | 1 |
| 18 | *CREBBP* | 8 | 4 | 0.368 | 1.488552 | 1 |
| 19 | *ANKRD1* | 3 | 1 | 0.4325 | 2.223996 | 1 |
| 20 | *CALR* | 3 | 1 | 0.4325 | 2.223996 | 1 |
| 21 | *NODAL* | 3 | 1 | 0.4325 | 2.223996 | 1 |
| 22 | *STRA6* | 3 | 1 | 0.4325 | 2.223996 | 1 |
| 23 | *ZEB2* | 3 | 1 | 0.4325 | 2.223996 | 1 |
| 24 | *LBR* | 4 | 2 | 0.4929 | 1.481908 | 1 |
| 25 | *RAI1* | 13 | 9 | 0.5315 | 1.069274 | 1 |
| 26 | *SMAD6* | 2 | 1 | 0.6123 | 1.478654 | 1 |
| 27 | *TAB2* | 2 | 1 | 0.6123 | 1.478654 | 1 |
| 28 | *TBX3* | 2 | 1 | 0.6123 | 1.478654 | 1 |
| 29 | *KCNH2* | 3 | 2 | 0.6396 | 1.108287 | 1 |
| 30 | *JAK2* | 5 | 4 | 0.6811 | 0.9216485 | 1 |
| 31 | *NOTCH1* | 6 | 5 | 0.6981 | 0.8837761 | 1 |
| 32 | *NSD1* | 6 | 5 | 0.6981 | 0.8837761 | 1 |
| 33 | *ELN* | 7 | 6 | 0.7134 | 0.8582199 | 1 |
| 34 | *MYH6* | 15 | 13 | 0.7378 | 0.844262 | 1 |
| 35 | *ALDH1A2* | 4 | 4 | 0.7875 | 0.7351475 | 1 |
| 36 | *NF1* | 4 | 4 | 0.7875 | 0.7351475 | 1 |
| 37 | *TBX5* | 2 | 2 | 0.7921 | 0.7369374 | 1 |

| | | | | | | |
|---|---|---|---|---|---|---|
| 38 | *PTCH1* | 7 | 7 | 0.8035 | 0.7324166 | 1 |
| 39 | *FOXH1* | 1 | 1 | 0.8201 | 0.7378233 | 1 |
| 40 | *LEFTY2* | 1 | 1 | 0.8201 | 0.7378233 | 1 |
| 41 | *RAF1* | 1 | 1 | 0.8201 | 0.7378233 | 1 |
| 42 | *TBX20* | 1 | 1 | 0.8201 | 0.7378233 | 1 |
| 43 | *NOS3* | 5 | 6 | 0.8697 | 0.6092539 | 1 |
| 44 | *MED13L* | 7 | 8 | 0.8701 | 0.6380351 | 1 |
| 45 | *MYH7* | 3 | 4 | 0.8794 | 0.549811 | 1 |
| 46 | *VEGFA* | 3 | 4 | 0.8794 | 0.549811 | 1 |
| 47 | *PDGFRA* | 2 | 3 | 0.8937 | 0.4894172 | 1 |
| 48 | *NPHP3* | 4 | 7 | 0.9589 | 0.4147024 | 1 |
| 49 | *GATA4* | 1 | 3 | 0.968 | 0.2441802 | 1 |
| 50 | *ZFPM2* | 2 | 5 | 0.9747 | 0.2912188 | 1 |
| 51 | *MYH11* | 0 | 6 | 1 | 0 | 1 |
| 52 | *NKX2-6* | 0 | 3 | 1 | 0 | 1 |
| 53 | *DLL1* | 0 | 2 | 1 | 0 | 1 |
| 54 | *EFNB2* | 0 | 2 | 1 | 0 | 1 |
| 55 | *F7* | 0 | 2 | 1 | 0 | 1 |
| 56 | *NOTCH2* | 0 | 2 | 1 | 0 | 1 |
| 57 | *SLC2A10* | 0 | 2 | 1 | 0 | 1 |

**Table S6. CHD patients carrying *SORBS2* variants.**

| Subjects | Cardiac Phenotypes* |
| --- | --- |
| P1 | TOF ASD PDA |
| P2 | PA VSD SV |
| P3 | DORV VSD PH |
| P4 | DORV CAVC VSD ASD PH |
| P5 | TOF ASD |
| P6 | TOF ASD PDA |
| P7 | DORV VSD PH |
| P8 | SA CAVC Dextrocardia PA |
| P9 | PA VSD ASD Dextrocardia |
| P10 | TOF ASD |
| P11 | TOF ASD |
| P12 | DORV VSD PH ASD |
| P13 | TA VSD ASD PH |
| P14 | TGA VSD PH |
| P15 | TGA VSD PDA ASD |
| P16 | TGA VSD ASD |
| P17 | DORV VSD ASD PH |
| P18 | TOF ASD |
| P19 | SV PA ASD |
| P20 | PA VSD |

*: TOF: Tetralogy of Fallot; ASD: Atrial septal defect; PDA: Patent ductus arteriosus; PA: Pulmonary atresia; VSD: Ventricular septal defect; SV: Single ventricle; DORV: Double outlet right ventricle; PH: Pulmonary hypertension; CAVC: complete atrioventricular septal defect; SA: Single atrium; TA: Tricuspid atresia; TGA: Transposition of great arteries.

**Table S7. Primers for qPCR.**

| Gene | Forward 5'- 3' | Reverse 5'- 3' |
|---|---|---|
| *OCT4* | GGGGTTCTATTTGGGAAGGT | CTGGTTCGCTTTCTCTTTCG |
| *NANOG* | CAGCCCCGATTCTTCCACCAGTCCC | CGGAAGATTCCCAGTCGGGTTCACC |
| *SOX2* | ATGACCAGCTCGCAGACCTA | GACTTGACCACCGAACCCAT |
| *T* | TATGAGCCTCGAATCCACATAGT | GTTCCTCCATCATCTCTTTGTGA |
| *MESP1* | TCGAAGTGGTTCCTTGGCAGAC | CCTCCTGCTTGCCTCAAAGTGTC |
| *TBX5* | AGACTCGCTGCTGAAAGGAC | GGAGCTGCACAGAATGTCAA |
| *HAND1* | AATCCTCTTCTCGACTGGGC | CCTTCAAGGCTGAACTCAAGA |
| *HCN4* | GACACCGCTATCAAAGTGGA | AGGTCCCAGTAAAATCTGAAGT |
| *ISL1* | TCACGAAGTCGTTCTTGCTG | CATGCTTTGTTAGGGATGGG |
| *MEF2C* | TTTCCTGTTTCCTCCAAACAA | CCAAGGACTAATCTGATCGGG |
| *TBX1* | CGGCTCCTACGACTATTGCCC | GGAACGTATTCCTTGCTTGCCCT |
| *NRARP* | CCAACTGCGAGTTCAACGTG | GGGAAGGTACAGCAGAGACG |
| *HEY1* | TGGTACCCAGTGCTTTTGAG | CTCCGATAGTCCATAGCAAGG |
| *HEYL* | ATGCAAGCCAGGAAGAAACGCAGA | AGCTTGGAAGAGCCCTGTTTCTCA |
| *PTCH1* | CCACAGAAGCGCTCCTACA | CTGTAATTTCGCCCCTTCC |
| *GLI1* | CTCCCGAAGGACAGGTATGTAAC | CCCTACTCTTTAGGCACTAGAGTT |
| *cTnT* | ATGAGCGGGAGAAGGAGCGGCAGAAC | TCAATGGCCAGCACCTTCCTCCTCTC |
| *MYH7* | ACCAACCTGTCCAAGTTCCG | TTCAAGCCCTTCGTGCCAAT |
| *MYH6* | TCCGTGAAGGGATAACCAGG | ACAGTCACCGTCTTCCCATTC |

| | | |
|---|---|---|
| *MLC-2A* | TCAAAGAAGCCTTCAGCTGTATC | TGAACTCATCCTTGTTCACCAC |
| *BMP4* | CGATGTGGGCTGGAATGA | TGGTTGAGTTGAGGTGGTCAG |
| *SMAD3* | CATCGAGCCCCAGAGCAATA | GTGGTTCATCTGGTGGTCACT |
| *ID2* | TGGACTCGCATCCCACTATT | CAGAAGCCTGCAAGGACAG |
| *MSX1* | CTCGTCAAAGCCGAGAGC | CGGTTCGTCTTGTGTTTGC |
| *FGF8* | TGAGCTGCCTGCTGTTGCACTT | TGAAGACGCAGTCCTTGCCTT |
| *FGF18* | GGACATGTGCAGGCTGGGCTA | GTAGAATTCCGTCTCCTTGCCCTT |
| *FZD5* | CTTGTTTCCAAAGTCCAATCAAGTG | GCCTACTCTTCACCCTTCTTTAACG |
| *FOXF1* | CAGCCGTATCTGCACCAGAA | ACTCCTTTCGGTCACACATGCT |

**Table S8. Subphenotypes of CHD cohort.**

| Cardiac Phenotypes | Subjects | Subjects Frequency (%) |
|---|---|---|
| Tetralogy of Fallot | 89 (46/43)* | 29.7 |
| Pulmonary atresia | 60 (19/41)* | 20 |
| Double outlet right ventricle | 59 (30/29)* | 19.7 |
| Transposition of great arteries | 46 (29/17)* | 15.3 |
| Single atrium/single ventricle | 27 | 9 |
| Tricuspid atresia | 11 | 3.7 |
| Interrupted aortic arch | 8 | 2.7 |
| **Total** | 300 | |

*, numbers within parentheses indicate the numbers of patients with/without ASD.

**Table S9. Known CHD genes included in targeted sequencing panel.**

| Gene | Phenotype | OMIM | Cytogenetic location | Genomic location(GRCh38) |
|---|---|---|---|---|
| *ACTC1* | ASD/VSD | 102540 | 15q14 | chr15: 34,788,095-34,795,725 |
| *ACVR1* | BAV/AVSD | 102576 | 2q24.1 | chr2: 157,736,445-157,875,895 |
| *ACVR2B* | Heterotaxy/PS/DORC/TGA | 602730 | 3p22.2 | chr3: 38,454,298-38,493,141 |
| *ALDH1A2* | TOF | 603687 | 15q21.3 | chr15: 57,953,423-58,065,922 |
| *ANKRD1* | TAPVC | 609599 | 10q23.31 | chr10: 90,912,099-90,921,274 |
| *ATRX* | Turner syndrome | 300032 | Xq21.1 | chrX: 77,504,877-77,786,268 |
| *BCOR* | Heterotaxy | 300485 | Xp11.4 | chrX: 40,051,245-40,177,389 |
| *BRAF* | Noonan/Cardiofaciocutaneous syndrome | 164757 | 7q34 | chr7: 140,719,326-140,924,927 |
| *CFC1* | Heterotaxy/TOF/TGA/IAA | 605194 | 2q21.1 | chr2: 130,592,164-130,599,574 |
| *CHD7* | Charge syndrome | 608892 | 8q12.2 | chr8: 60,678,743-60,868,027 |
| *CITED2* | ASD/VSD | 602937 | 6q24.1 | chr6: 139,372,254-139,374,649 |
| *COL2A1* | Stickler syndrome/AVSD | 120140 | 12q13.11 | chr12: 47,972,964-48,006,211 |
| *CREBBP* | Rubinstein-Taybi syndrome/ ventricular septation | 600140 | 16p13.3 | chr16: 3,725,053-3,880,726 |
| *CRELD1* | AVSD/ASD | 607170 | 3p25.3 | chr3: 9,933,787-9,945,412 |
| *CSDE1* | 1p13.2 microdeletion | 191510 | 1p13.2 | chr1: 114,716,912-114,758,049 |
| *EHMT1* | Kleefstra syndrome | 607001 | 9q34.3 | chr9: 137,618,991-137,836,126 |
| *ELN* | PPS/SVAS/AS | 130160 | 7q11.23 | chr7: 74,027,771-74,069,906 |

| | | | | |
|---|---|---|---|---|
| *EVC* | Ellis-van Creveld syndrome | 604831 | 4p16.2 | chr4: 5,711,196-5,829,042 |
| *EVC2* | Ellis-van Creveld syndrome | 607261 | 4p16.2 | chr4: 5,562,407-5,709,547 |
| *FBN1* | BAV/Marfan syndrome | 134797 | 15q21.1 | chr15: 48,408,305-48,645,787 |
| *FLNA* | ASD | 300017 | Xq28 | chrX: 154,348,531-154,374,637 |
| *FOXC1* | TOF/Aortic valve dysplasia | 601090 | 6p25.3 | chr6: 1,610,445-1,613,896 |
| *FOXH1* | TOF/TGA/ASD/IAA | 603621 | 8q24.3 | chr8: 144,473,731-144,476,334 |
| *FOXL2* | VSD | 605597 | 3q22.3 | chr3: 138,944,223-138,947,139 |
| *G6PC3* | ASD/PVS/PDA | 611045 | 17q21.31 | chr17: 44,070,699-44,076,343 |
| *GATA4* | ASD/AVSD/TOF/VSD | 600576 | 8p23.1 | chr8: 11,676,918-11,760,001 |
| *GATA6* | ASD/PTA/PS/TOF | 601656 | 18q11.2 | chr18: 22,169,436-22,202,527 |
| *GDF1* | DORV/Heterotaxy/TGA/TOF | 602880 | 19p13.11 | chr19: 18,868,545-18,896,143 |
| *GJA1* | AVSD/HLHS | 121014 | 6q22.31 | chr6: 121,435,576-121,449,743 |
| *GPC3* | DORV | 300037 | Xq26.2 | chrX: 133,535,744-133,985,645 |
| *HAND2* | VSD/PS/TOF | 602407 | 4q34.1 | chr4: 173,526,500-173,530,226 |
| *HEY2* | TOF | 604674 | 6q22.31 | chr6: 125,747,638-125,762,242 |
| *HOXA1* | VSD | 142955 | 7p15.2 | chr7: 27,092,992-27,096,005 |
| *HRAS* | Costello syndrome/ASD/VSD | 190020 | 11p15.5 | chr11: 532,241-535,566 |
| *IRX4* | VSD | 606199 | 5p15.33 | chr5: 1,877,426-1,887,178 |
| *JAG1* | Alagille syndrome/TOF | 601920 | 20p12.2 | chr20: 10,637,683-10,674,045 |
| *KMT2D* | HLHS/VSD/ASD | 602113 | 12q13.12 | chr12: 49,018,974-49,060,883 |
| *KRAS* | Noonan/Cardiofaciocutaneous syndrome | 190070 | 12p12.1 | chr12: 25,204,788-25,251,002 |

| | | | | |
|---|---|---|---|---|
| *LBR* | Human CHD Gene | 600024 | 1q42.12 | chr1: 225,401,501-225,428,854 |
| *LEFTY2* | Heterotaxy/TGA/AVSD | 601877 | 1q42.12 | chr1: 225,936,597-225,941,491 |
| *MAP2K1* | Cardiofaciocutaneous syndrome | 176872 | 15q22.31 | chr15:66,386,872-66,491,543 |
| *MAP2K2* | Cardiofaciocutaneous syndrome | 601263 | 19p13.3 | chr19:4,090,320-4,124,183 |
| *MED13L* | TGA | 608771 | 12q24.21 | chr12: 115,958,575-116,277,218 |
| *MGP* | Keutel syndrome | 154870 | 12p12.3 | chr12: 14,880,863-14,886,061 |
| *MID1* | Opitz syndrome | 300552 | Xp22.2 | chrX: 10,445,309-10,833,689 |
| *MYH11* | PDA/TAAD | 160745 | 16p13.11 | chr16: 15,703,134-15,857,031 |
| *MYH6* | ASD/Tricuspid atresia | 160710 | 14q11.2 | chr14: 23,381,989-23,408,276 |
| *MYH7* | Ebstein's anomaly of tricuspid valve/ASD | 160760 | 14q11.2 | chr14: 23,412,737-23,435,685 |
| *NF1* | TOF | 613113 | 17q11.2 | chr17: 31,094,926-31,377,676 |
| *NKX2-5* | ASD/HLHS/TOF/VSD | 600584 | 5q35.1 | chr5: 173,232,103-173,235,320 |
| *NKX2-6* | PTA/VSD/TOF/DORV | 611770 | 8p21.2 | chr8: 23,702,450-23,706,597 |
| *NODAL* | Heterotaxy/TOF | 601265 | 10q22.1 | chr10: 70,431,935-70,447,947 |
| *NOTCH1* | BAV/LVOTO/TOF | 190198 | 9q34.3 | chr9: 136,494,432-136,545,785 |
| *NOTCH2* | Alagille syndrome | 600275 | 1p12 | chr1: 119,911,552-120,069,702 |
| *NPHP3* | ASD/PTA/PDA | 608002 | 3q22.1 | chr3: 132,680,608-132,722,458 |
| *NRAS* | Noonan syndrome | 164790 | 1p13.2 | chr1: 114,704,463-114,716,893 |
| *NSD1* | Sotos syndrome | 606681 | 5q35.3 | chr5: 177,131,834-177,300,212 |
| *PDGFRA* | TAPVC | 173490 | 4q12 | chr4: 54,229,088-54,298,246 |
| *PTPN11* | Noonan/LEOPARD syndrome | 176876 | 12q24.13 | chr12: 112,418,897-112,509,917 |

| | | | | |
|---|---|---|---|---|
| *RAF1* | Noonan/LEOPARD syndrome | 164760 | 3p25.2 | chr3: 12,583,600-12,664,200 |
| *RAI1* | Smith-Magenis syndrome | 607642 | 17p11.2 | chr17: 17,681,375-17,811,452 |
| *RBM10* | TARP syndrome | 300080 | Xp11.3 | chrX: 47,145,195-47,186,814 |
| *ROR2* | Robinow syndrome | 602337 | 9q22.31 | chr9: 91,722,595-91,950,205 |
| *SALL4* | Duane-radial ray syndrome/VSD | 607343 | 20q13.2 | chr20: 51,782,716-51,802,522 |
| *SH3PXD2B* | Frank-Ter Haar syndrome | 613293 | 5q35.1 | chr5: 172,325,180-172,454,522 |
| *SHOC2* | Noonan syndrome | 602775 | 10q25.2 | chr10: 110,919,369-111,013,666 |
| *SLC2A10* | Arterial tortuosity syndrome | 606145 | 20q13.12 | chr20: 46,708,357-46,736,346 |
| *SMAD6* | BAV/AS/COA | 602931 | 15q22.31 | chr15: 66,702,109-66,781,999 |
| *SOS1* | Noonan syndrome | 182530 | 2p22.1 | chr2: 38,981,548-39,124,958 |
| *STRA6* | PDAC syndrome | 610745 | 15q24.1 | chr15: 74,179,465-74,212,266 |
| *TAB2* | VSD/BAV/LVOTO | 605101 | 6q25.1 | chr6: 149,217,923-149,411,612 |
| *TBX1* | DiGeorge syndrome/TOF | 602054 | 22q11.21 | chr22: 19,756,702-19,783,592 |
| *TBX20* | TOF/PTA/ASD/VSD | 606061 | 7p14.2 | chr7: 35,199,935-35,254,099 |
| *TBX3* | Ulnar-mammary syndrome | 601621 | 12q24.21 | chr12: 114,670,253-114,684,163 |
| *TBX5* | Holt-Oram syndrome | 601620 | 12q24.21 | chr12: 114,353,910-114,408,707 |
| *TDGF1* | TOF/VSD | 187395 | 3p21.31 | chr3: 46,574,554-46,582,462 |
| *TFAP2B* | PDA/PTA/TOF | 601601 | 6p12.3 | chr6: 50,817,691-50,847,618 |
| *VEGFA* | TOF | 192240 | 6p21.1 | chr6: 43,770,208-43,786,486 |
| *ZEB2* | Mowat-Wilson syndrome | 605802 | 2q22.3 | chr2: 144,384,374-144,520,390 |
| *ZFPM2* | TOF | 603693 | 8q23.1 | chr8: 105,318,858-105,804,538 |
| *ZIC3* | Heterotaxy/TGA/ASD/PS | 300265 | Xq26.3 | chrX: 137,566,126-137,577,690 |

**Table S10. Candidate CHD genes included in targeted sequencing panel.**

| Gene | OMIM | Cytogenetic location | Genomic location(GRCh38) |
|---|---|---|---|
| *CALR* | 109091 | 19p13.13 | chr19: 12,938,599-12,944,489 |
| *CNN1* | 600806 | 19p13.2 | chr19: 11,538,716-11,550,322 |
| *CRK* | 164762 | 17p13.3 | chr17: 1,421,352-1,456,266 |
| *DLL1* | 606582 | 6q27 | chr6: 170,282,199-170,290,608 |
| *EFNB2* | 600527 | 13q33.3 | chr13: 106,489,730-106,535,039 |
| *EPOR* | 133171 | 19p13.2 | chr19: 11,377,204-11,384,341 |
| *ETS1* | 164720 | 11q24.3 | chr11: 128,458,760-128,587,592 |
| *F7* | 613878 | 13q34 | chr13: 113,105,772-113,120,680 |
| *FLI1* | 193067 | 11q24.3 | chr11: 128,685,262-128,813,266 |
| *GJA5* | 121013 | 1q21.2 | chr1: 147,756,198-147,781,126 |
| *HEY1* | 602953 | 8q21.13 | chr8: 79,764,010-79,767,863 |
| *JAK2* | 147796 | 9p24.1 | chr9: 4,985,085-5,128,182 |
| *KCNH2* | 152427 | 7q36.1 | chr7: 150,944,955-150,978,313 |
| *MYCN* | 164840 | 2p24.3 | chr2: 15,940,437-15,947,006 |
| *NFATC1* | 600489 | 18q23 | chr18: 79,395,771-79,529,322 |
| *NOS3* | 163729 | 7q36.1 | chr7: 150,991,055-151,014,598 |
| *PDLIM3* | 605889 | 4q35.1 | chr4: 185,500,659-185,535,557 |
| *PTCH1* | 601309 | 9q22.32 | chr9: 95,442,979-95,517,056 |
| *QKI* | 609590 | 6q26 | chr6: 163,414,485-163,578,595 |

| | | | |
|---|---|---|---|
| *RPS6KA2* | 601685 | 6q27 | chr6: 166,409,363-166,862,550 |
| *SHH* | 600725 | 7q36.3 | chr7: 155,799,983-155,812,272 |
| *SLC25A4* | 103220 | 4q35.1 | chr4: 185,143,262-185,150,383 |
| *SORBS2* | 616349 | 4q35.1 | chr4: 185,585,443-185,956,715 |
